# Complex-valued representations of time-series gene expression profiles for network analysis

**DOI:** 10.64898/2026.06.16.732574

**Authors:** Jianqiang Sun, Wei Cao, Ikumi Kamachi, Kentaro K. Shimizu, Jun Sese

## Abstract

Time-series RNA sequencing provides a powerful framework for studying dynamic gene regulation, yet conventional analyses usually represent gene expression profiles as real-valued vectors in Euclidean space and quantify similarity using correlation or distance. Inspired by quantum information theory, we present a framework for encoding time-series gene expression profiles as complex-valued vectors comprising amplitude and phase components in Hilbert space. We designed multiple encoding models to represent gene expression in the amplitude of complex-valued vectors, encode temporal differences in the phase, and extend the phase representation to incorporate the direction of local expression changes. Gene-gene similarity was then quantified using fidelity, which measures the overlap between two encoded vectors. Evaluation using time-series RNA-seq datasets across diverse species and biological contexts showed that different encoding models produced distinct fidelity distributions that were related to, but distinct from, conventional correlation measures. We then constructed gene-gene networks using pairwise fidelity values and detected communities containing genes with similar temporal profiles. Although fidelity distributions differed across encoding models, the resulting communities captured major temporal expression programs, and functional annotations based on gene ontology and Kyoto encyclopedia of genes and genomes pathway analyses provided exploratory biological context. The detected communities were comparable to those obtained using conventional methods, including weighted correlation network analysis and fuzzy c-means clustering. Furthermore, as a proof-of-concept, we performed SWAP-test circuit simulations to mimic fidelity computation on a quantum computer; under noise-aware conditions, these simulations produced less accurate fidelity estimates with higher computational cost than classical computation. As a proof-of-concept, this study provides a complementary view of temporal transcriptome organization, rather than a uniformly superior alternative to conventional methods.

## Introduction

High-throughput RNA sequencing (RNA-seq) enables genome wide quantification of gene expression across diverse biological contexts. Time-series RNA-seq experiments capture temporal trajectories of gene activity and provide a basis for studying dynamic regulatory processes underlying development, environmental responses, cellular differentiation, and physiological maturation across diverse biological systems [1–5]. A central task in the analysis of such data is to quantify similarity between temporal expression profiles, which supports clustering, gene co-expression network construction, and the identification of functionally related gene groups. Conventional approaches usually represent each gene as a real-valued vector in Euclidean space and evaluate similarity using correlation- or distance-based measures. For example, weighted gene co-expression network analysis (WGCNA) [6] is widely used to detect correlation-based gene modules, whereas fuzzy c-means clustering (FCM) [7] provides a distance-based approach for grouping genes with similar expression profiles.

Concepts from quantum information theory provide a mathematical framework in which data can be represented as complex-valued state vectors in a Hilbert space [8]. In this setting, each state vector consists of amplitude and phase components, where amplitudes encode magnitudes and phases define relative angular relationships, yielding a geometric representation of the data. Time-series expression profiles provide a natural setting to explore such a representation because expression level and temporal changes between adjacent time points can be assigned to distinct components of the same vector. For example, temporal changes can be represented as phase shifts, so that temporal progression is described as angular variation in the complex plane. Similarity between state vectors can then be quantified by their overlap, defined as the squared magnitude of the inner product and commonly referred to as fidelity [8]. Fidelity ranges from 0 to 1, with higher values indicating stronger geometric alignment between state vectors. In this framework, fidelity reflects both amplitude alignment and phase coherence, allowing gene–gene similarity to be defined in terms of Hilbert-space geometry rather than only pointwise agreement in Euclidean space. Although this formalism originates from quantum physics, it can also be interpreted more generally as a structured geometric embedding of data [9, 10]. Its potential utility as a representation framework for temporal gene expression dynamics, however, remains largely unexplored.

Here, we represent time-series gene expression profiles using state vectors, in which each gene is described by amplitude and phase components across temporal basis states. Gene–gene similarity is quantified using fidelity, allowing similarity relationships to be interpreted as geometric overlap in Hilbert space rather than as pointwise agreement in Euclidean space. Using real time-series RNA-seq datasets, we examine how this representation affects pairwise similarity relationships and downstream fidelity-based structures. Although formally related to concepts from quantum information theory, the proposed framework is intended as a mathematical representation rather than a physical quantum model. It therefore serves as a proof-of-concept for reinterpreting temporal gene expression dynamics in a geometric setting.

## Materials and Methods

### RNA-seq datasets and expression quantification

Five time-series RNA-seq datasets were used in this study to evaluate the framework across diverse biological contexts: *Arabidopsis thaliana* (Col-0), wheat (*Triticum aestivum*, strain Shiluan02-1), soybean (*Glycine max*, Williams 82), human (*Homo sapiens*, cell line WA09), and mouse (*Mus musculus*, strain CD-1). Raw FASTQ data for all datasets were obtained from the National Center for Biotechnology Information Sequence Read Archive.

The Arabidopsis dataset consisted of shoot apical meristem samples collected daily from 7 to 16 days after germination (DAG), with two biological replicates per time point [1]. The wheat dataset consisted of leaf samples collected at eight weekly time points spanning the pre-vernalization, vernalization, and post-vernalization phases, with three biological replicates per time point [2]. The soybean dataset consisted of leaf and cotyledon samples collected at five developmental stages, beginning at the four leaf stage and then 2, 4, 6, and 7 weeks later during the reproductive stage, with three biological replicates per time point [3]. In this study, we only used leaf samples. The human dataset was derived from an in vitro differentiation time course of human embryonic stem cells into cortical neurons over 77 days [4]. From the original experiment, we used only samples cultured in cortical differentiation medium under the 90% confluency condition, comprising six unevenly spaced time points at days 19, 26, 33, 49, 63, and 77 of differentiation, with two to three biological replicates per time point. The mouse dataset was derived from the study of mammalian organs across different developmental stages [5]. From the original experiments across multiple species and organs, we used only mouse postnatal heart samples, comprising five unevenly spaced time points at postnatal days 0, 3, 14, 28, and 63, with three to four biological replicates per time point.

All datasets were processed using the same analysis pipeline. Raw sequencing reads were subjected to quality filtering using Trimmomatic v0.40 [11]. Cleaned reads were aligned using STAR v2.7.11b [12] to the TAIR10 reference genome for Arabidopsis [13], IWGSC reference genome v2 for wheat [14], Wm82.a2.v1 reference genome for soybean [15], and the GENCODE GRCh38 and GRCm39 reference genomes and corresponding gene annotations for human and mouse, respectively [16]. Gene-level read counts were quantified using featureCounts v2.1.1 [17] with the corresponding gene annotation files for each reference genome. The count data were then normalized using the variance stabilizing transformation implemented in DESeq2 v1.48.1 [18].

### Preprocessing of temporal expression profiles

For each dataset, genes were retained if their variance-stabilized expression value exceeded 5 in at least one-third of the libraries. Among the retained genes, temporally variable genes were selected using mean absolute deviation thresholds of 0.5 for Arabidopsis, human, and mouse and 0.8 for wheat and soybean. The higher threshold for wheat and soybean was used because these species have substantially larger gene repertoires than the other datasets, largely owing to polyploidy or whole-genome duplication events and the subsequent retention of duplicated gene copies. Biological replicates were then averaged at each time point to generate an expression matrix with genes as rows and time points as columns for downstream analyses. This averaging step was used because the primary aim of this study was to provide a proof-of-concept evaluation of encoding strategies for temporal expression profiles, rather than to construct a formal model of variation among biological replicates. The sensitivity of fidelity estimates and community detection to variation among replicates was assessed separately by resampling biological replicate samples (Supplementary Note S1).

Let *t=*1,2,…,*T* index the ordered sampling points in each dataset, where *T* denotes the number of observed time points rather than the final sampling time. Then, let *x*_*g,t*_ ≥0 denote the expression magnitude, that is, the variance-stabilized expression value, of gene *g* at time point *t*. The temporal expression profile of gene *g* was represented as

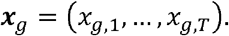

For each gene, expression profiles were standardized across time points using z-score normalization,

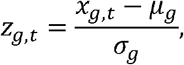

where *µ*_*g*_ and *σ*_*g*_ denote the mean and standard deviation of across time points, respectively.

To encode temporal changes, we computed the interval averaged expression magnitude between adjacent time points,

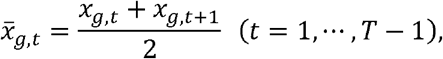

and the temporal difference of the standardized expression profile,

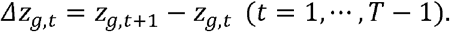

Because some datasets contained unevenly spaced time points, we defined interval weights to account for differences in sampling interval length. Let *τ*_*t*_ denote the actual sampling time associated with index *t*; thus *τ*_*T*_ corresponds to the final sampling time. Then, define the adjacent interval as

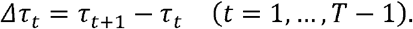

The normalized interval weight was defined as

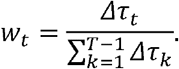

This weight represents the relative duration of each adjacent time interval within the full time series.

### Complex-valued encoding of temporal expression profiles

Temporal expression profiles were encoded as normalized complex-valued state vectors. The main concept of the encodings was to assign expression magnitude to amplitude components and temporal changes to phase components. We considered five encodings: expression amplitude encoding (EA), temporal difference phase encoding (TDP), integrated difference phase encoding (IDP), orthogonal-direction temporal difference phase encoding (ODTDP), and orthogonal-direction integrated difference phase encoding (ODIDP).

The EA encoding model was designed to encode gene expression magnitudes in the amplitudes of a state vector without incorporating temporal changes. In this model, each gene *g* was encoded as

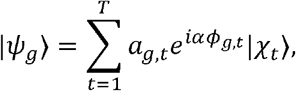

where 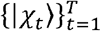 denotes an orthonormal temporal basis. The amplitude was defined as 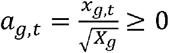, where 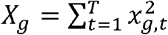. The definition satisfies 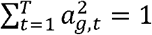, thereby ensuring that ~ *ψ*_*g*_ ⟩ is normalized (i.e, ⟨ *ψ*_*g*_ ~ *ψ*_*g*_ ⟩=1). The phase was defined as *ϕ*_*g,t*_ =*t*. Because this phase term is common across genes, it does not introduce gene-wise temporal changes into the EA model. Thus, EA serves as a baseline encoding in which gene–gene similarity is determined by normalized expression magnitudes. The parameter *α* controls phase scaling. Unless otherwise stated, the default value was set to 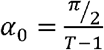.

To incorporate temporal changes into the state-vector representation, we defined two phase-based encodings: TDP and IDP. In both models, the basis dimension is *T* −1, corresponding to temporal intervals between adjacent time points. Each basis state ~ *χ*_*t*_ ⟩ represents the interval between *τ*_*t*_ and *τ*_*t*+1_. Each gene was encoded as

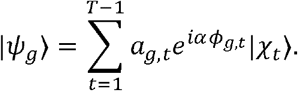

The amplitude was defined as 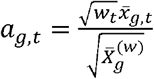, where 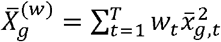 to ensure 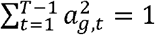. Thus, the amplitude component represents expression magnitude on temporal intervals under the elapsed time measure. In the TDP model, the phase was defined as *ϕ*_*g,t*_=Δ*Z*_*g,t*_,which represents similarity of local temporal changes. The default phase-scaling parameter was set to 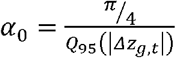, where *Q*_95_ (·) denotes the 95th percentile computed over all genes and temporal intervals in each dataset. In the IDP model, the phase was defined as 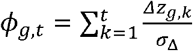, where *σ*_Δ_ denotes the standard deviation of Δ*Z*_*g,t*_ computed across all genes and temporal intervals. This cumulative term is equivalent to the displacement from the initial standardized expression value after scaling by *σ*_Δ_, because 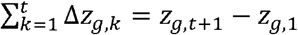. Therefore, the IDP represents cumulative trajectory similarity relative to the initial time point. The default scaling parameter was set to 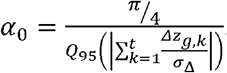.

We further extended the TDP and IDP models to encode the direction of expression change using orthogonal-direction bases. These models are referred to as ODTDP and ODIDP, respectively. For each temporal interval *t*, two orthogonal basis states, ~*χ*_*t*,+_⟩and, ~*χ*_*t*,−_ were introduced to represent increasing and decreasing expression changes. Each gene was encoded as

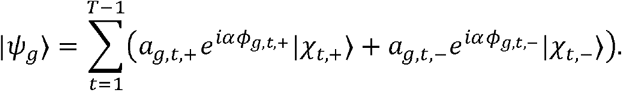

The direction weights were defined as 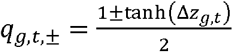. The corresponding amplitudes were defined as 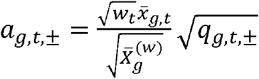. Because *q*_*g,t*+_ + *q*_*g,t*−_ =1, these amplitudes satisfy the normalization condition over the expanded orthogonal basis, 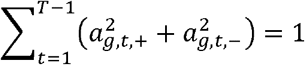. The phase definitions were inherited from the corresponding phase-based encodings. For the ODTDP, *ϕ*_*g,t*±_=Δ*Z*_*g,t*_, whereas for the ODIDP, 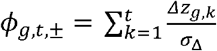. The default scaling parameter *α*_0_ for each orthogonal-direction model was set to the same value as that used in its corresponding phase-based model.

### Calculation of fidelity

Similarity between genes *g* and *h* was quantified using quantum fidelity,

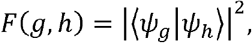

where ~ *ψ*_*g*_ ⟩ and ~ *ψ*_*h*_ ⟩ denote the normalized state vectors of genes *g* and *h*, respectively. Because all state vectors were normalized, fidelity values ranged from 0 to 1.

Fidelity values were computed analytically from the state vectors using NumPy v2.4.1, and this analytical calculation was used for all main analyses. For visualization and correlation analyses, 100,000 off-diagonal gene– gene pairs were randomly sampled without replacement from the upper triangle of the pairwise similarity matrix for each dataset. The same set of sampled pairs was used consistently across all models.

As a supplementary proof-of-concept, SWAP-test quantum circuits were implemented using Qiskit v2.3.0 [19] and simulated on classical computers to examine whether the analytically computed fidelity values could be reproduced in a circuit-based formulation (Supplementary Note S2 and S3). All experiments were conducted with Python 3.11.13 on an Ubuntu 24.04 system equipped with an Intel Core i9-11900K processor (8 cores, 16 threads, 3.5 GHz).

### Construction of temporal similarity networks and community detection

For each dataset and encoding model, the pairwise fidelity matrix was converted into a weighted, undirected network, where nodes represent genes and edge weights correspond to fidelity values. Self-connections were excluded. Edges were filtered by applying a fidelity threshold and a mutual k-nearest-neighbor criterion, retaining only gene pairs that satisfied both conditions. Communities, defined here as groups of genes that are more densely connected to one another than to the rest of the network, were detected using the Leiden algorithm [20]. Communities containing fewer than 20 genes were excluded from downstream analyses.

To determine appropriate network configurations, three hyperparameters were explored by grid search: the number of nearest neighbors, the fidelity cutoff, and the Leiden resolution parameter (see Supplementary Method S1 for details). For each parameter combination, network construction and Leiden community detection were repeated 100 times to account for algorithmic stochasticity. For each configuration, modularity, assigned gene count, community count, and median community size were summarized across runs, and partition stability was quantified as the mean pairwise adjusted mutual information (AMI) across the 100 partitions. Here, assigned gene count refers to the total number of input genes assigned to any detected community, community count refers to the number of detected communities, and community size refers to the number of genes in each community. Configurations were first filtered to exclude unstable, weakly structured, or excessively fragmented solutions according to predefined criteria for AMI, modularity, assigned gene count, community count, and median community size. Among the retained configurations, Pareto-optimal solutions were identified by maximizing modularity, AMI, assigned gene count, and median community size while minimizing community count. From the Pareto front, a final parameter set was selected as a balanced solution by minimizing the distance to the ideal point after min–max normalization of each objective.

To evaluate the sensitivity of network construction to the phase-scaling parameter, fidelity matrices were also computed over a range of scaling parameters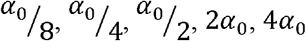, and 8*α*_0_. To separate the effect of *α* from downstream graph-construction choices, network parameters were selected only at the default phase scale (i.e., *α= α*_0_) for each encoding model. The selected values of *k* and the Leiden resolution parameter were then fixed and applied unchanged across all *α* values.

### Conventional approach benchmarks

To benchmark the fidelity-based networks, we also constructed conventional co-expression networks and detected gene modules using WGCNA v1.73 [6], and performed FCM clustering of gene expression profiles using Mfuzz v2.68.0 [21]. For WGCNA, network construction followed the standard workflow implemented in the blockwiseModules function using variance-stabilized expression values. Signed weighted networks were constructed using a signed adjacency matrix and a signed topological overlap matrix. Gene modules were identified by hierarchical clustering based on topological overlap matrix dissimilarity followed by dynamic tree cutting, with a minimum module size of 20 genes. The soft thresholding power was selected as the smallest value that achieved a scale free topology fit index greater than 0.8. The selected powers were 28 for Arabidopsis and 29 for wheat. For the soybean, human, and mouse datasets, powers greater than 30 were required to satisfy this criterion. Because the software implementation restricted the maximum tested power to 30, a power of 30 was used for these datasets.

For Mfuzz, z-scored variance-stabilized expression values were used as input. The optimal number of clusters was selected by examining the decrease in the minimum centroid distance as the number of clusters increased and choosing the point at which further increases yielded only marginal separation gains. The selected numbers of clusters were 7, 15, 7, 9, and 5 for Arabidopsis, wheat, soybean, human, and mouse, respectively.

Note that differential time-course methods such as maSigPro and ImpulseDE2 were considered but were not included as primary benchmarks. These methods are designed primarily to identify genes showing differential temporal behavior between experimental conditions or treatment groups. Because the objective of this study was to define and evaluate unsupervised gene–gene similarity structures within individual temporal expression datasets, WGCNA and Mfuzz were selected as more appropriate conventional comparators.

### Functional annotation over-representation analysis

Gene ontology (GO) over-representation analysis was performed for each detected community from each encoding model using clusterProfiler v4.16.0 [22]. GO annotations were obtained from TAIR database for Arabidopsis, Supplementary Table S2 of Nomura et al for wheat [23], JGI Data Portal for soybean, and annotation packages of org.Hs.eg.db v3.21.0 and org.Mm.eg.db v3.21.0 for human and mouse. Analyses were conducted separately for each dataset and were restricted to the biological process category. GO term enrichment was evaluated using a hypergeometric test, with the genes retained for network construction used as the background gene universe. Because this analysis was intended to provide exploratory functional annotation of gene sets in community levels rather than formal statistical validation of the detected communities or encoding models, GO terms with raw p-values < 0.05 were retained as nominally enriched candidate terms.

For Arabidopsis and mouse datasets, Kyoto encyclopedia of genes and genomes (KEGG) pathway over-representation analysis was additionally performed using clusterProfiler with the KEGG gene annotations obtained through Bioconductor annotation resources. KEGG pathway enrichment was evaluated using the same background gene sets and statistical framework as the GO analysis.

## Results

### Fidelity distributions reflect encoding design and data structure

In this framework, gene expression magnitudes and temporal changes were encoded as the amplitude and phase components, respectively, of normalized state vectors for each gene (Fig. 1A, B). Similarity between genes was then quantified using fidelity for each encoding model. The distribution of fidelity values differed substantially across encoding strategies and datasets (Fig. 1C; Supplementary Table S1).

**Figure 1.**
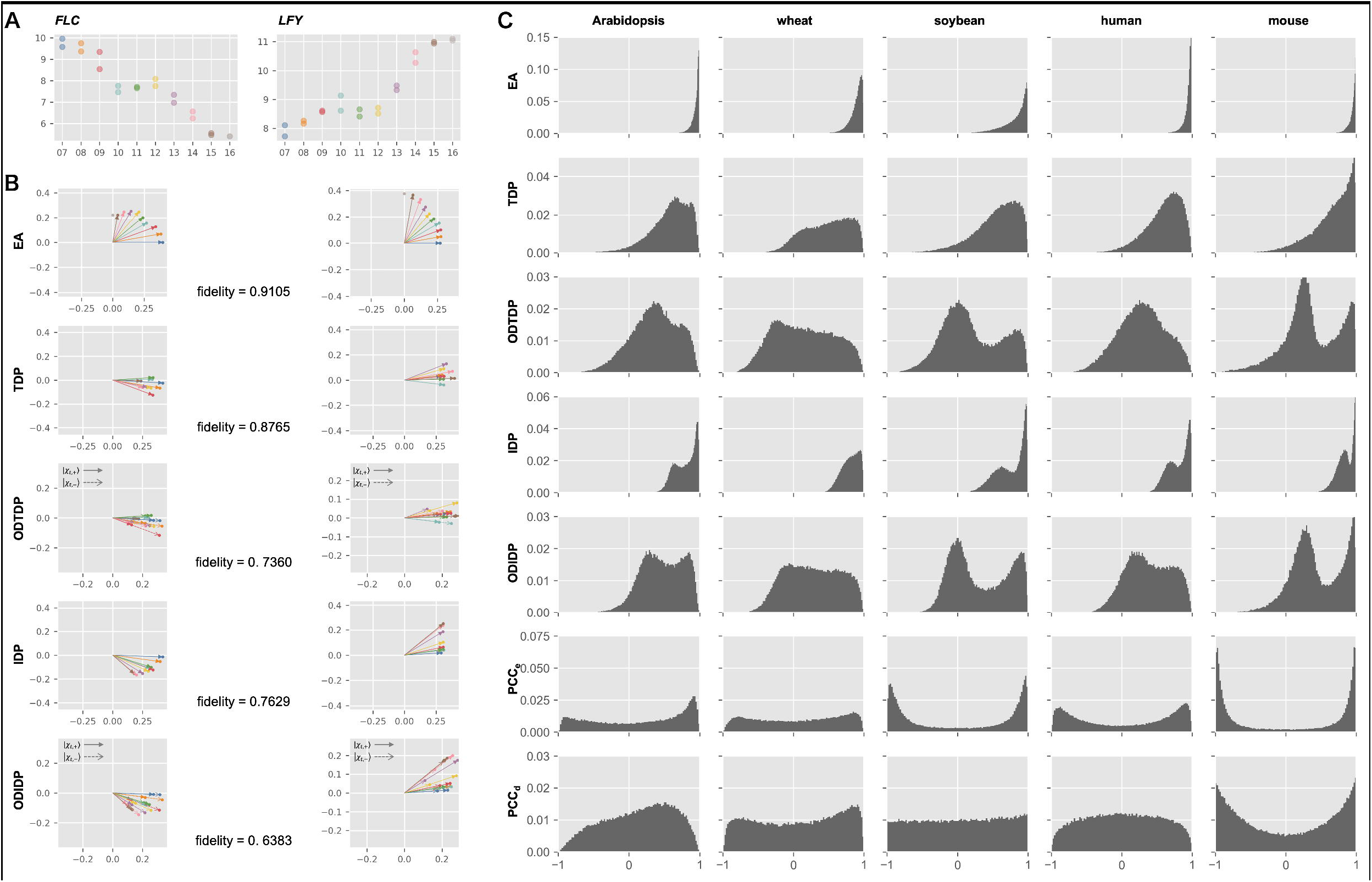
Visualization of complex-valued encodings and distributions of pairwise similarity values. (A) Time-series expression profiles of *FLC* and *LFY* in the Arabidopsis dataset. The x-axis indicates days after germination, and the y-axis indicates normalized expression values. (B) Complex-plane visualization of the encoded vectors for *FLC* and *LFY* under EA, TDP, ODTDP, IDP, and ODIDP models. Each arrow represents one complex-valued component associated with a temporal basis or temporal-interval basis. The fidelity between FLC and LFY is shown for each encoding model. In ODTDP and ODIDP, each temporal interval is decomposed into two orthogonal direction components representing increasing and decreasing local expression changes. Solid arrows indicate increasing-direction components, and dotted arrows indicate decreasing-direction components. (C) Histograms of pairwise fidelity values for each encoding model and Pearson correlation coefficients computed from expression profiles (PCC_e_) and temporal differences (PCC_d_). For each dataset, distributions were computed using 100,000 randomly sampled off-diagonal gene pairs.

Across the datasets, EA produced the narrowest fidelity distributions, with values concentrated near the upper end of the range. In EA, the phase term is shared across genes and cancels in pairwise fidelity, reducing the measure to squared cosine similarity between normalized expression magnitude profiles (Supplementary Note S4). In addition, because these profiles contain nonnegative values, their inner products do not involve cancellation by negative components, which can shift EA fidelity values toward high values.

The phase-based models, TDP and IDP, produced broader distributions than EA, but their distributional shapes differed according to the phase definition and dataset. TDP, which encodes local temporal changes in the phase component, generally broadened the fidelity distribution relative to EA. IDP, which represents expression state relative to the initial time point, often showed a narrower distribution than TDP and tended to show a more apparent two-component pattern. In these cases, one component was concentrated near high fidelity values, whereas another appeared at lower fidelity values with a smaller peak.

The orthogonal-direction models, ODTDP and ODIDP, generally produced broader distributions than their corresponding phase-based models and modified their peak structures. This tendency can be understood from their basis design, in which each temporal interval is divided into positive and negative direction components according to the sign and magnitude of the local temporal difference. Gene pairs with concordant local change directions retain overlap within the same direction components, whereas pairs with opposite local change directions have reduced overlap because their contributions are assigned to different components.

The tendency toward two-component distributions in IDP and orthogonal-direction models appeared to depend on the interaction between encoding design and dataset structure. When a dataset contains strongly polarized increasing and decreasing trajectories, IDP can separate gene pairs according to their cumulative displacement from the initial standardized expression value. Gene pairs moving in similar directions from the initial state tend to retain high fidelity, whereas pairs moving in divergent directions can form a lower fidelity component. Similarly, orthogonal-direction models can make two-component patterns more apparent by separating same direction and opposite direction local changes into different components. This behavior was more evident in datasets such as soybean and mouse, but it was not observed uniformly across all temporal expression structures.

### Pairwise model comparisons reveal concordance and absolute differences

Pairwise comparisons of fidelity values between encoding models showed that the similarity structure differed among models and datasets (Table 1; Supplementary Fig. S1). Among comparisons involving EA, IDP showed the highest Pearson correlation coefficients (PCC) with EA across all five datasets. This pattern is consistent with the encoding design of IDP. Although IDP is defined by accumulating temporal differences, the accumulated value at each point corresponds to the change from the initial standardized expression value. In pairwise fidelity, the initial component contributes a global phase that is specific to each gene pair and does not affect the squared modulus. Consequently, IDP is more closely aligned with similarity based on expression state than TDP is.

**Table 1.**
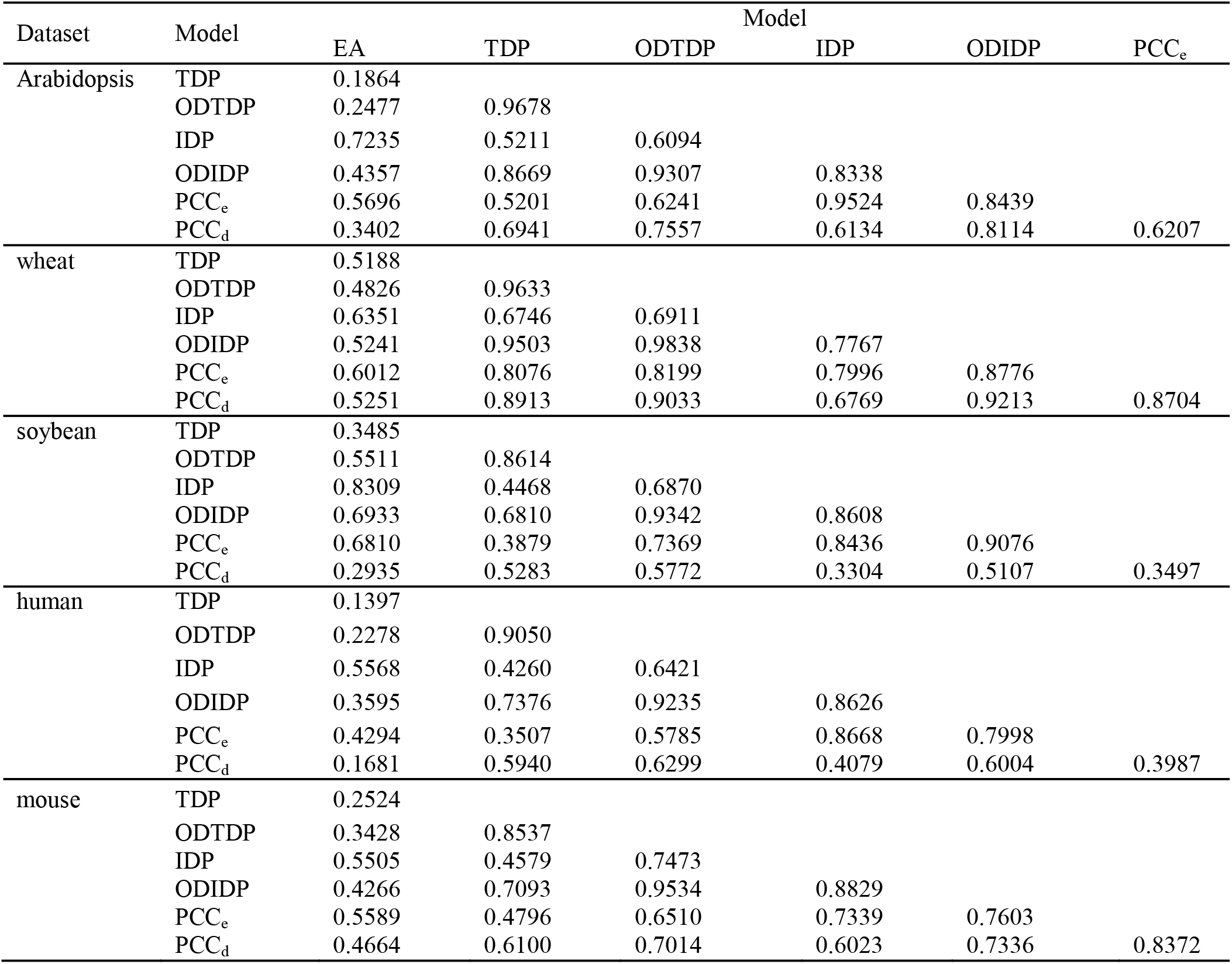
Pairwise correlations between encoding models and reference measures. Values represent Pearson correlation coefficients between the pairwise similarity values obtained from the models or reference measures shown in the corresponding row and column. PCC_e_ and PCC_d_ denote Pearson correlation coefficients computed from expression profiles and temporal differences, respectively.

| Dataset | Model | Model |  |  |  |  |  |
|---|---|---|---|---|---|---|---|
|  |  | EA | TDP | ODTDP | IDP | ODIDP | PCC<sub>e</sub> |
| Arabidopsis | TDP | 0.1864 |  |  |  |  |  |
|  | ODTDP | 0.2477 | 0.9678 |  |  |  |  |
|  | IDP | 0.7235 | 0.5211 | 0.6094 |  |  |  |
|  | ODIDP | 0.4357 | 0.8669 | 0.9307 | 0.8338 |  |  |
|  | PCC<sub>e</sub> | 0.5696 | 0.5201 | 0.6241 | 0.9524 | 0.8439 |  |
|  | PCC<sub>d</sub> | 0.3402 | 0.6941 | 0.7557 | 0.6134 | 0.8114 | 0.6207 |
| wheat | TDP | 0.5188 |  |  |  |  |  |
|  | ODTDP | 0.4826 | 0.9633 |  |  |  |  |
|  | IDP | 0.6351 | 0.6746 | 0.6911 |  |  |  |
|  | ODIDP | 0.5241 | 0.9503 | 0.9838 | 0.7767 |  |  |
|  | PCC<sub>e</sub> | 0.6012 | 0.8076 | 0.8199 | 0.7996 | 0.8776 |  |
|  | PCC<sub>d</sub> | 0.5251 | 0.8913 | 0.9033 | 0.6769 | 0.9213 | 0.8704 |
| soybean | TDP | 0.3485 |  |  |  |  |  |
|  | ODTDP | 0.5511 | 0.8614 |  |  |  |  |
|  | IDP | 0.8309 | 0.4468 | 0.6870 |  |  |  |
|  | ODIDP | 0.6933 | 0.6810 | 0.9342 | 0.8608 |  |  |
|  | PCC<sub>e</sub> | 0.6810 | 0.3879 | 0.7369 | 0.8436 | 0.9076 |  |
|  | PCC<sub>d</sub> | 0.2935 | 0.5283 | 0.5772 | 0.3304 | 0.5107 | 0.3497 |
| human | TDP | 0.1397 |  |  |  |  |  |
|  | ODTDP | 0.2278 | 0.9050 |  |  |  |  |
|  | IDP | 0.5568 | 0.4260 | 0.6421 |  |  |  |
|  | ODIDP | 0.3595 | 0.7376 | 0.9235 | 0.8626 |  |  |
|  | PCC<sub>e</sub> | 0.4294 | 0.3507 | 0.5785 | 0.8668 | 0.7998 |  |
|  | PCC<sub>d</sub> | 0.1681 | 0.5940 | 0.6299 | 0.4079 | 0.6004 | 0.3987 |
| mouse | TDP | 0.2524 |  |  |  |  |  |
|  | ODTDP | 0.3428 | 0.8537 |  |  |  |  |
|  | IDP | 0.5505 | 0.4579 | 0.7473 |  |  |  |
|  | ODIDP | 0.4266 | 0.7093 | 0.9534 | 0.8829 |  |  |
|  | PCC<sub>e</sub> | 0.5589 | 0.4796 | 0.6510 | 0.7339 | 0.7603 |  |
|  | PCC<sub>d</sub> | 0.4664 | 0.6100 | 0.7014 | 0.6023 | 0.7336 | 0.8372 |

TDP and IDP showed moderate correlations across datasets, indicating that local temporal change similarity and cumulative state similarity capture related but distinct aspects of temporal expression profiles. In contrast, ODTDP and ODIDP showed strong correlations across datasets. This high concordance is consistent with their shared orthogonal separation of positive and negative local changes. In both ODTDP and ODIDP, gene pairs with opposite local change directions are assigned to different direction components and therefore have reduced overlap, even though the two models use different phase definitions.

The orthogonal-direction extension affected the TDP and IDP formulations differently. In general, TDP remained more strongly correlated with ODTDP than IDP did with ODIDP. This pattern is consistent with the model definitions: both the TDP phase and the direction components in ODTDP are based on local temporal differences, whereas IDP represents cumulative state relative to the initial time point and ODIDP adds local direction information to this cumulative phase representation. Thus, the orthogonal-direction extension tended to preserve the broad similarity structure of TDP more strongly than that of IDP. However, this tendency was not universal, with exceptions observed in datasets such as soybean and mouse, indicating that the effect of orthogonal-direction encoding also depends on data structure.

The mean absolute error (MAE) provided an additional perspective on these relationships (Supplementary Table S2). Even when model pairs were strongly correlated, their absolute fidelity values were not always identical. In particular, the orthogonal-direction extension produced larger absolute changes when applied to IDP than when applied to TDP across datasets, even when the corresponding correlations remained high.

### Fidelity captures similarity patterns distinct from Pearson correlation

We additionally compared fidelity with Pearson correlation coefficients computed from expression profiles (PCC_e_) and from temporal difference profiles (PCC_d_), as reference measures rather than as benchmarks or optimization targets. The distributions of PCC_e_ and PCC_d_ were dataset dependent (Fig. 1C; Supplementary Table S1). PCC_e_ showed varying degrees of positive and negative polarization across datasets, whereas PCC_d_ often showed distinct distributional shapes. This indicates that similarity of expression states and similarity of local temporal changes were not equivalent.

These distributional differences helped explain the model dependent relationship between fidelity and PCC (Table 1). TDP and ODTDP generally showed stronger correlations with PCC_d_ than with PCC_e_, consistent with their use of local temporal differences in the phase component. In contrast, IDP and ODIDP generally showed stronger correspondence with PCC_e_, consistent with the structure of IDP, in which the accumulated temporal difference corresponds to expression state relative to the initial time point.

The distributional shapes of PCC_e_ and PCC_d_ also helped explain when two component patterns appeared in fidelity distributions. When PCC_e_ showed sharp peaks near both positive and negative extremes, IDP tended to show a clearer two component pattern, with gene pairs separating according to whether their trajectories moved in similar or divergent directions from the initial state. Orthogonal-direction models could also show two-component patterns when local change directions were strongly polarized, although this tendency depended on the dataset and encoding model. Similarly, TDP distributions were more closely related to PCC_d_. When PCC_d_ was strongly concentrated near the correlation extremes, TDP tended to show a sharp high-fidelity peak, whereas datasets with less polarized PCC_d_ distributions showed smoother TDP distributions with broader peaks.

Despite these relationships, fidelity was not equivalent to conventional PCC. In the TDP and IDP models, a subset of gene pairs showed high fidelity despite strong negative PCC_e_ values, using the criterion of fidelity greater than 0.8 and PCC_e_ less than −0.8 (Supplementary Fig. S2; Supplementary Table S3). These discordant high-fidelity cases were mainly associated with oppositely directed monotonic changes in TDP (Supplementary Fig. S2A). This can occur because TDP depends on coherence among local phase differences rather than the global direction of the expression trajectory (Supplementary Note S5). In IDP, they involved large, oppositely directed initial changes followed by nearly stable or mutually matched expression levels (Supplementary Fig. S2B), reflecting the cumulative phase definition in which stable offsets after the initial discrepancy can preserve phase coherence (Supplementary Note S6). The orthogonal-direction models strongly reduced these directionally inconsistent assignments. Under the same criterion, ODTDP reduced such pairs to negligible levels across datasets, and ODIDP showed zero or near zero counts (Supplementary Table S3). These results indicate that fidelity defines encoding-dependent similarity structures that are related to, but distinct from, conventional correlation measures.

### Encoding model affects temporal similarity network resolution

Fidelity values from each encoding model were used to construct weighted gene networks, followed by community detection to identify groups of genes with high pairwise fidelity. The resolution and composition of the detected communities varied among encoding models and datasets (Fig. 2; Supplementary Figs. S3–S7). Adding temporal phase information and separating local change directions altered the degree of community subdivision, but the extent of this effect depended on the data structure. Despite these differences in partitioning resolution, the major temporal trends were detected across encoding models.

**Figure 2.**
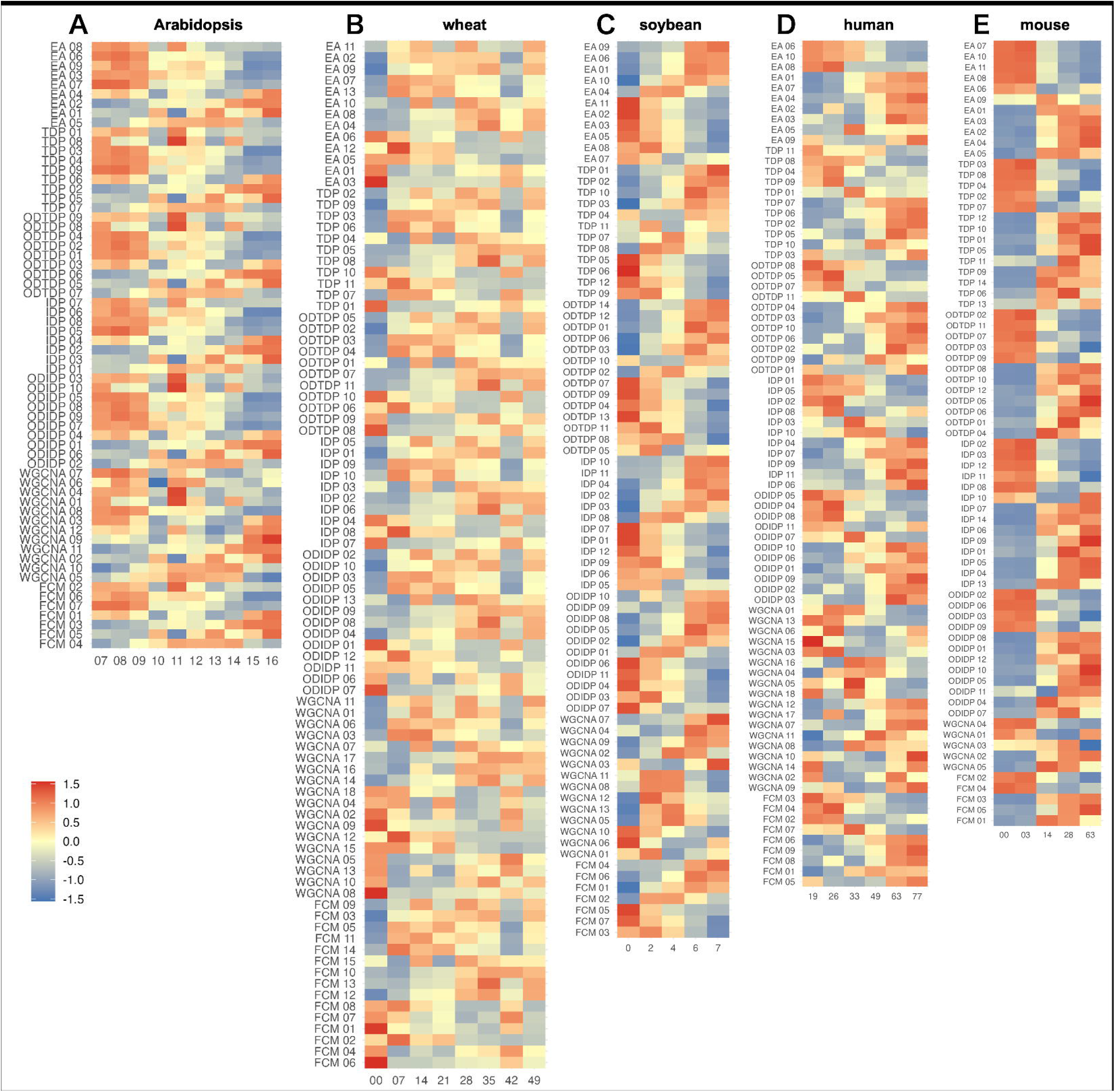
Average gene expression profiles within detected communities. Heatmaps show the average z-scored expression profiles of genes within communities detected by the five encoding models (EA, TDP, ODTDP, IDP, and ODIDP), WGCNA, and fuzzy c-means (FCM) clustering. Rows indicate communities labeled by model name and community numbers, and columns indicate dataset-specific time points. The color scale represents the average z-scored expression value within each community at each time point. Results are shown for (A) Arabidopsis, (B) wheat, (C) soybean, (D) human, and (E) mouse datasets.

Inspection of community expression profiles revealed exceptions specific to individual models. In TDP, some communities contained genes with broadly opposite overall trajectories, whereas in IDP some communities contained genes with opposite initial changes followed by nearly stable or mutually similar later profiles. For example, in the human dataset, TDP Communities 02 (TDP-02), TDP-11, IDP-08, and IDP-11 showed such patterns (Supplementary Fig. S6). These patterns are consistent with the pairwise analysis showing that discordant gene pairs can receive high-fidelity values in TDP and IDP despite low PCC values (Supplementary Fig. S2; Supplementary Table S3; Supplementary Notes S5 and S6). Such discordant patterns were reduced in the orthogonal-direction models, although they were not completely eliminated.

Comparison with WGCNA and FCM further highlighted differences in community resolution. In Arabidopsis, soybean, and human, WGCNA and the fidelity-based models generally produced more resolved partitions than FCM, whereas in wheat the relative resolution was more comparable among methods. Mouse showed a distinct pattern: both WGCNA and FCM produced only five modules or clusters, markedly fewer than in the other datasets. In this dataset, WGCNA and FCM grouped the data into a small number of broad increasing or decreasing programs, whereas the fidelity-based models provided a finer subdivision of these broad temporal trends. Across datasets, the average expression profiles of WGCNA modules and FCM clusters overlapped qualitatively with those detected by the fidelity-based models, indicating that the main temporal programs were shared across methods despite differences in partitioning resolution.

### Functional annotations provide biological context for fidelity-based communities

In Arabidopsis, the detected communities included transient profiles around 10–11 DAG (e.g., EA-01, EA-08, TDP-01, TDP-05, ODTDP-03, ODTDP-05, IDP-03, IDP-07, ODIDP-04, ODIDP-06), broadly increasing profiles (e.g., EA-02, TDP-02, ODTDP-06, IDP-02, ODIDP-01), and broadly decreasing profiles (e.g., EA-03, TDP-03, ODTDP-04, IDP-05, ODIDP-05) across models (Fig. 2A; Supplementary Fig. S3). Over-representation analyses of GO and KEGG pathway annotations supported the interpretation that these communities captured developmental transition together with accompanying metabolic, hormonal, and stress-related programs during shoot apical meristem development (Supplementary Data S1 and S2). Specifically, transient communities were linked to growth and reproductive organ development (GO:0099402, GO:0061458), suggesting a transition from vegetative growth toward reproductive development. Both increasing and decreasing communities were associated with glucosinolate-related processes and stress responses (GO:0019760, GO:0006950). In addition, although KEGG pathway annotations are biased toward metabolic pathways, the over-represented pathways supported a phase transition involving hormone signaling, photosynthetic activity, translation, and carbohydrate metabolism in selected communities (KEGG:ath04075, KEGG:ath00196, KEGG:ath03010, KEGG:ath00500).

In wheat, the detected communities represented multiple temporal profiles across the pre-vernalization, vernalization, and post-vernalization phases, including early-increasing profiles (e.g., TDP-03, ODTDP-03, IDP-09, ODIDP-03), early-decreasing profiles (e.g., TDP-01, ODTDP-08, IDP-07, ODIDP-07), and transient profiles at early (e.g., TDP-04, ODTDP-01, IDP-03, ODIDP-06) and late time points (e.g., TDP-08, TDP-09, ODTDP-10, ODTDP-11, IDP-01, IDP-06, ODIDP-01, ODIDP-04) across multiple models (Fig. 2B; Supplementary Fig. S4). Functional annotations suggested that the wheat communities reflected several broad aspects of the vernalization time course rather than a single cold-response trajectory (Supplementary Data S3). In detail, early-increasing communities were linked to reproductive developmental growth (GO:0061458, GO:0048507), whereas early-decreasing communities were associated with multiple biological processes, including stress responses such as cold response (GO:0009409), photosynthesis (GO:0015979), and hormone responses involving abscisic acid and ethylene (GO:0009738, GO:0009873). Communities with early transient were associated with stress responses, including cold and hypoxia responses, as well as growth-related processes (GO:0009409, GO:0001666, GO:0007049, GO:1905393, GO:0048527). By contrast, communities with late transient were associated with photosynthesis, reproductive and developmental processes, and proteolysis (GO:0015979, GO:0022414, GO:0048507, GO:0006508).

In soybean, the detected community profiles were relatively simple despite the larger number of communities, including increasing, decreasing, and transient profiles (Fig. 2C; Supplementary Fig. S5). GO annotations suggested that these communities mainly reflected changes in leaf physiological state during plant development, with signals related to photosynthetic function, carbohydrate metabolism, regulatory activity, lipid metabolism, transport, and stress responses (Supplementary Data S4).

In human cortical differentiation, the detected communities contained increasing, decreasing, and transient profiles (Fig. 2D; Supplementary Fig. S6). GO annotations of these communities included biological processes related to proliferative activity, cellular specialization, and structural organization during the time course. These annotations reflected broad cellular programs accompanying in vitro differentiation and were consistent with the experimental design (Supplementary Data S5).

In mouse postnatal heart development, the detected communities were dominated by monotonic increasing profiles (e.g., TDP-01, IDP-04) or monotonic decreasing profiles (e.g., TDP-03, IDP-02), with additional transient profiles around postnatal days 14–28 (e.g., TDP-02, TDP-06, IDP-08, IDP-13) (Fig. 2E; Supplementary Fig. S7). GO and KEGG pathway annotations suggested that these communities reflected broad postnatal heart maturation programs (Supplementary Data S6 and S7). Specifically, late-increasing or transient communities were associated with proliferative and chromosome-related programs, suggesting developmental changes in cell-cycle activity during postnatal maturation (GO:0007049, GO:0007059, KEGG:mmu04110). Other communities reflected broader tissue remodeling and state-transition processes, including heart or muscle development (GO:0003007, GO:0030198, KEGG:mmu04510, KEGG:mmu05415).

Overall, functional annotations supported the biological interpretability of the fidelity-based communities and were broadly consistent with the biological settings of the five time-series datasets. However, they should be regarded as exploratory context rather than formal validation of the community structures.

### Sensitivity to variation among biological replicates

We also examined whether the use of expression values averaged across biological replicates affected fidelity-based network structures. Fidelity values computed from expression values averaged across replicates and from resampled expression matrices showed strong agreement across datasets and encoding models (PCC = 0.886– 0.999; MAE = 0.002–0.059; Supplementary Note S1; Supplementary Fig. S8). Networks reconstructed from resampled mean fidelity matrices retained largely similar sets of assigned genes, although exact community assignments showed moderate agreement with those obtained in the main analysis (Supplementary Table S4). The average temporal expression profiles of communities reconstructed from resampled mean fidelity matrices were nevertheless broadly similar to those observed in the main analysis (Supplementary Fig. S9). These results indicate that variation among biological replicates can affect community boundaries and exact gene membership, while the broader temporal programs recovered by the networks are qualitatively preserved.

### Phase scaling modulates fidelity distributions

The phase assigned to each component of a state vector determines its complex orientation. Therefore, changing the phase scaling parameter *α* is expected to alter the overlap between state vectors. We varied *α* to examine how phase scaling affected fidelity distributions. As expected from the shared phase term in EA, the EA model was unaffected by phase scaling, whereas the phase-based and orthogonal-direction models showed systematic changes in their fidelity distributions across datasets (Supplementary Figs. S10). Smaller phase scales generally compressed fidelity values toward the upper end of the range, although the extent of this compression differed among models and datasets. As *α* increased, the distributions generally broadened and shifted toward lower fidelity values. At larger phase scales, such as 4*α*_0_ and 8*α*_0_, many distributions showed increased density at lower fidelity values, consistent with stronger phase dispersion and reduced overlap between state vectors. These results indicate that *α* modulates similarity geometry through its effect on phase.

We next examined whether these changes in fidelity distributions affected network construction and community detection. For each encoding model, networks were reconstructed across *α* values using the network parameters selected at the baseline phase scale, *α* = *α*_0_. As a result, community numbers and community size distributions remained broadly similar across *α* values within the same model, although some model and dataset specific deviations were observed at extreme phase scales (Fig. 3). However, larger scales (e.g., ≥ 4 *α*_0_) increased within community heterogeneity in some phase-based models, where more discordant expression profiles were grouped into the same communities (Supplementary Fig. S11). These discordant patterns were reduced in the orthogonal-direction models, although they were not completely eliminated.

**Figure 3.**
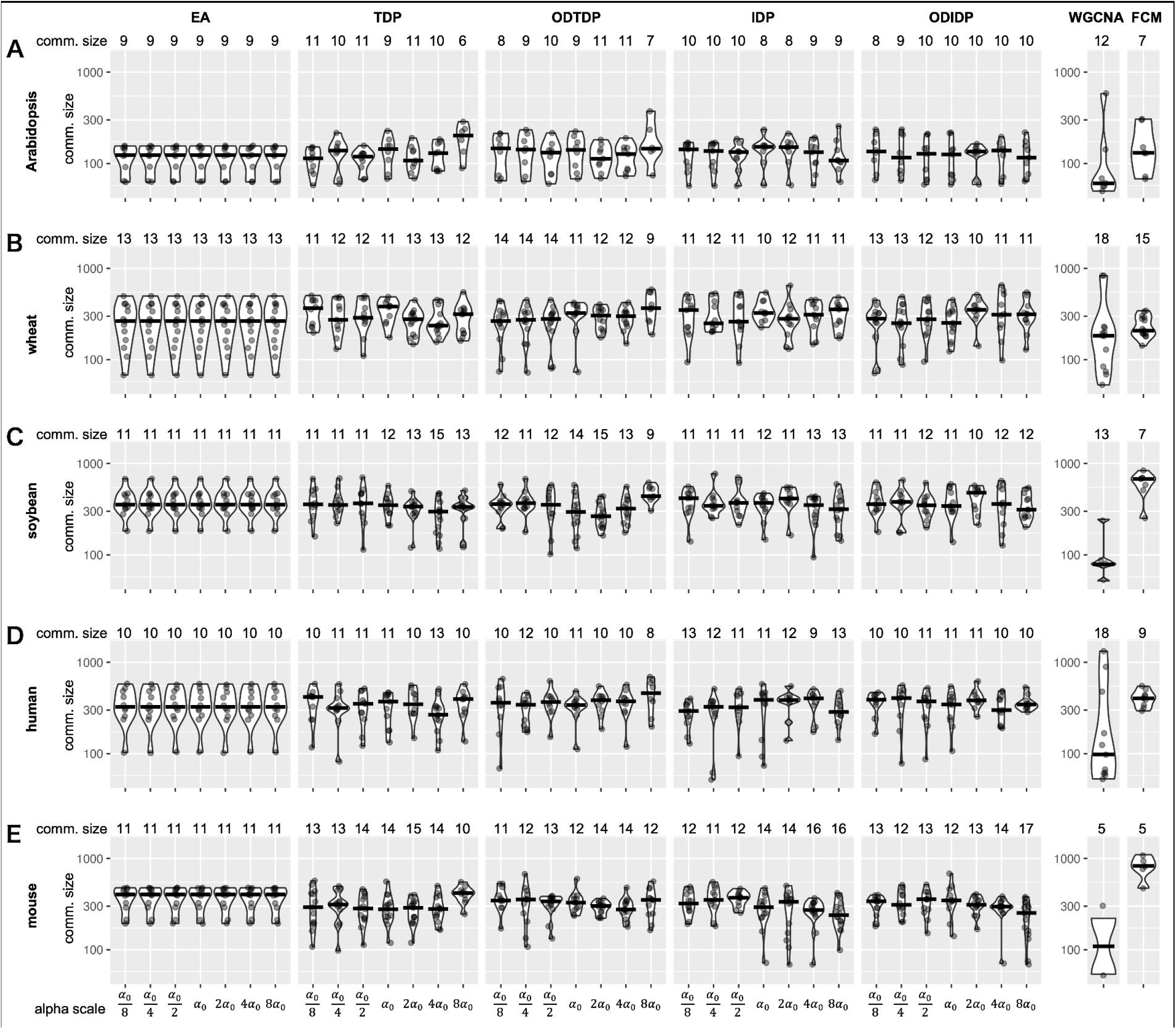
Community number and community size distributions across models and phase scales. Violin plots show the distribution of community sizes, defined as the number of genes per detected community, across encoding models and phase-scaling values for (A) Arabidopsis, (B) wheat, (C) soybean, (D) human, and (E) mouse datasets. For the fidelity-based models, community detection was evaluated across phase scales from 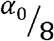 to 8*α*_0_, where *α*_0_ denotes the default phase-scaling parameter for each encoding model. WGCNA and fuzzy c-means (FCM) clustering are shown as conventional comparison methods. Each point represents one detected community, and black horizontal bars indicate median community size. Numbers above the violins indicate the number of detected communities under each condition.

### Circuit simulation reproduces analytical trends but remains less accurate

To assess computational consistency, fidelity was additionally estimated using quantum circuit simulations based on the SWAP test (Supplementary Note S2). These simulations were performed on classical computers to emulate quantum circuit execution. Although analytical and circuit estimates showed high correlations, with values up to 0.99, substantial discrepancies remained, with root mean square errors on the order of 10^−1^ across models. These results indicate that, under the simulation setting used here, circuit estimates reproduced the overall analytical trends but were less accurate than direct analytical computation. Together with the higher computational cost, this limits the practical applicability of circuit simulation for large scale pairwise gene similarity analysis in the present setting.

## Discussion

We present this study as a proof-of-concept for representing time-series gene expression profiles using complex-valued state vectors and quantifying gene–gene similarity using fidelity, rather than as an attempt to establish fidelity as a universally superior alternative to conventional similarity measures. In this framework, amplitude components encode expression magnitude; in phase-based and orthogonal-direction encodings, elapsed sampling intervals are incorporated through amplitude weighting, so that the amplitude contribution reflects both expression level and interval duration. Phase components determine how temporal changes influence the overlap between state vectors. Fidelity is defined as the squared magnitude of the inner product between normalized state vectors, providing a geometric measure of overlap in Hilbert space.

Encoding design affects fidelity distributions and downstream community resolution because it determines how expression magnitude, temporal phase, and basis assignment contribute to geometric overlap. For example, in TDP, gene pairs with opposite overall expression trajectories can still receive high fidelity when their encoded components remain coherent in Hilbert space. This illustrates that encoding design should be guided by the definition of fidelity and by the temporal relationships that should be treated as similar or separated. Therefore, the practical goal should not be to identify a single encoding that is optimal for all time-series datasets. Instead, encodings should be designed according to the biological question, experimental design, and temporal relationship of interest. For example, when the objective is to identify genes with different initial expression levels or early transitions but similar later trajectories, extensions of IDP may be appropriate because they can emphasize coherence of cumulative expression states rather than pointwise agreement across the entire profile.

The phase-scaling parameter *α* should be treated as part of model specification and sensitivity analysis, rather than as a fixed universal constant. In the datasets examined here, phase scales up to 2*α*_*0*_ yielded qualitatively similar fidelity distributions and community resolutions, whereas larger *α* values tended to increase discordant high-fidelity gene pairs with distinct expression profiles and heterogeneity within communities. Thus, both encoding choice and *α* tuning should be guided by the temporal structure of the dataset and the biological question being addressed.

Fidelity-based communities captured major expression trends in the analyzed datasets, but similar broad trends were also captured by WGCNA and FCM. This qualitative agreement supports the view that the methods recover related temporal programs, whereas differences in community subdivision indicate that they emphasize different aspects of the same datasets. Because there is no external ground truth for the correct community resolution in these unsupervised analyses, differences among methods should be interpreted as reflecting different views of temporal expression organization rather than as evidence that one approach is uniformly preferable.

Several limitations should be noted. First, the encodings examined here represent only a limited subset of possible complex-valued feature maps, and future work should design encodings according to the experimental design and biological objective. Second, community detection depends not only on fidelity values but also on network construction parameters. Detected communities should therefore be interpreted as useful summaries of parameterized fidelity-based similarity networks rather than as uniquely determined biological modules. Third, functional annotations are biased toward model organisms and well-characterized metabolic or signaling pathways, and should be interpreted as supplementary biological context rather than formal validation of the encoding models or detected communities. Finally, the circuit-based implementation remains a conceptual demonstration. The present analyses rely on analytical fidelity computation by classical linear algebra and do not establish a quantum computational advantage.

## Availability of data and materials

The raw RNA-seq datasets used in the case studies were downloaded from the NCBI SRA database under the following accession nos.: PRJNA268115 for Arabidopsis, PRJNA963171 for wheat, PRJNA262564 for soybean, PRJNA244622 for human, and PRJEB26869 for mouse dataset. Functions for quantum state encoding and fidelity computation are implemented with Python and available on GitHub repository https://github.com/bitdessin/xqubit. The processed expression data, quantified from the RNA-seq datasets, along with analysis scripts supporting the findings of this study, are available on Zenodo under the identifier doi:10.5281/zenodo.20438943.

## Acknowledgements

Computations were partially performed on the SHIHO supercomputer at National Agriculture and Food Research Organization. ChatGPT, developed by OpenAI, was utilized to assist with proofreading and enhance the clarity of manuscript.

## Supplementary information

Supplementary_file: A PDF file containing Supplementary Method S1, Notes S1–S6, Figures S1–S11.

Supplementary_tables: An Excel file containing Supplementary Tables S1–S4.

Supplementary_data: A ZIP archive containing supplementary data in Excel format (Data S1–S7).

