## Supplemental Data 2 for "Complex-valued representations of time-series gene expression profiles for network analysis"

### Supplemental Materials

#### Supplementary Methods

##### Supplementary Method S1: Hyperparameter optimization for fidelity-based network construction

To identify appropriate network configurations for each dataset and encoding strategy, we performed a systematic grid search over three hyperparameters: the number of nearest neighbors ($k$), the fidelity cutoff, and the Leiden resolution parameter. Network parameters were selected at the default phase-scaling parameter for each encoding model.

Fidelity-based edge filtering was applied using two complementary types of cutoffs. Absolute fidelity thresholds of 0.7, 0.8, 0.9, 0.99, and 0.999 were considered, together with percentile-based thresholds corresponding to the 60th, 70th, 80th, 85th, 90th, and 95th percentiles of the upper-triangular elements of the fidelity matrix. The number of nearest neighbors (k) was varied across 10, 20, 25, 30, 35, 40, and 50. The Leiden resolution parameter was tested at 0.8, 1.0, 1.5, and 2.0.

For each combination of hyperparameters, network construction and community detection were repeated 100 times to account for stochasticity in the Leiden algorithm. The resulting partitions were evaluated using five metrics: (i) modularity; (ii) stability, defined as the mean adjusted mutual information (AMI) across all pairwise comparisons of the 100 partitions; (iii) assigned gene count, defined as the total number of input genes assigned to any detected community; (iv) community count, defined as the number of detected communities; and (v) median community size, defined as the median number of genes per community. To exclude unstable, weakly structured, or excessively fragmented solutions, parameter combinations were filtered according to predefined criteria. Specifically, only configurations with mean modularity > 0.4, mean AMI > 0.7, mean assigned gene count > 1,000, mean community count ≤ 21, and median community size > 20 were retained for further consideration.

Among the retained configurations, Pareto-optimal solutions were identified in a five-objective space. The mean modularity, mean AMI, mean assigned gene count, and median community size were treated as objectives to be maximized, whereas mean community count was treated as an objective to be minimized to avoid excessive fragmentation. For final selection from the Pareto front, each objective was transformed by min–max normalization across the retained configurations. For objectives to be maximized, the normalized score was defined as $s = \frac{x - x_{min}}{x_{max} - x_{min}}$, whereas for community count, which was minimized, the normalized score was defined as $s = \frac{x_{max} - x}{x_{max} - x_{min}}$. Thus, larger normalized scores consistently represented more favorable solutions for all objectives. The ideal point was defined as the point at which all normalized objective scores were equal to 1. From the Pareto front, the final parameter set was selected as the balanced solution with the smallest distance to this ideal point.

#### Supplementary Notes

##### Supplementary Note S1: Sensitivity of fidelity-based networks to variation among biological replicates

To assess the sensitivity of fidelity estimates and network structures to variation among biological replicates, 1,000 resampled gene-by-timepoint matrices were generated for each dataset. In each resampling iteration, one biological replicate sample was randomly selected within each time point, independently across time points, and the complete expression vector of the selected sample was used for all genes at the corresponding time point. Each resampled matrix was processed using the same standardization, encoding, and fidelity calculation procedures as in the main analysis, yielding one fidelity matrix per iteration. The resulting fidelity matrices were averaged elementwise across the 1,000 iterations to obtain a resampled mean fidelity matrix for each dataset and encoding model. Networks and communities were then reconstructed from this matrix using the same network construction procedure and the same hyperparameters selected in the main analysis using expression values averaged across replicates, without re-optimization.

Pairwise fidelity values computed from expression values averaged across replicates and from resampled expression matrices showed strong agreement across datasets and encoding models, with Pearson correlation coefficients (PCC) ranging from 0.886 to 0.999 and mean absolute error (MAE) ranging from 0.002 to 0.059 (Supplementary Fig. S8). EA and IDP showed particularly close agreement in most datasets, whereas relatively larger deviations were observed for TDP and orthogonal-direction (ODTDP and ODIDP) encodings, especially in soybean and mouse. These results indicate that pairwise fidelity values were broadly preserved after resampling of biological replicates, although encodings based on local temporal differences and direction-separated components were more sensitive to variation among replicates.

Communities were then compared between networks reconstructed from expression values averaged across replicates and those reconstructed from resampled mean fidelity matrices. Agreement was quantified using the Jaccard index of assigned gene sets and adjusted mutual information (AMI), with adjusted Rand index (ARI), normalized mutual information (NMI), and variation of information (VI) reported as complementary metrics. Metrics were computed both for all input genes, with unassigned genes treated as a single category, and for genes assigned to communities in both analyses. The sets of genes assigned to retained communities were largely preserved, with Jaccard indices ranging from 0.849 to 0.997 across all comparisons (Supplementary Table S5). In contrast, exact gene assignments to specific communities showed more moderate agreement. AMI values ranged from 0.458 to 0.820 when all input genes were considered and from 0.494 to 0.825 when the comparison was restricted to genes assigned to communities in both analyses (Supplementary Table S5).

Agreement was highest in Arabidopsis and generally lower in soybean, mouse, and some human models. Temporal-difference-based and orthogonal-direction encodings tended to show lower agreement than EA or IDP in several datasets, consistent with their stronger dependence on local temporal changes and direction-separated components. Despite these differences in exact gene membership, the average temporal expression profiles of communities reconstructed from resampled mean fidelity matrices were broadly similar to those observed in the main analysis (Supplementary Fig. S9). Major increasing, decreasing, transient, and biphasic temporal patterns were retained across datasets and encoding models.

Together, these results indicate that variation among biological replicates had limited effects on the overall pairwise fidelity structure and on the set of genes retained in communities, but could affect network thresholding, community boundaries, and exact gene assignment. A likely explanation is that multiple communities can share similar temporal patterns, so modest changes in fidelity values can alter the allocation of boundary genes among related communities during mutual nearest-neighbor filtering and Leiden community detection. Thus, the broader temporal programs recovered by the networks were qualitatively preserved, whereas gene-level community membership should be interpreted with some uncertainty near community boundaries.

#### Supplementary Note S2: Validation of fidelity estimation using quantum circuit simulations

To validate the computational implementation of state-vector-based similarity, we implemented the SWAP test using Qiskit and compared circuit-based fidelity estimates with analytically computed fidelity values obtained using NumPy. The number of qubits required for state preparation depends on the dimensionality of the encoded state vectors, which is determined by the number of time points and, for orthogonal-direction encodings, by the expansion of temporal intervals into direction-specific basis components (see Supplementary Note S3 for examples).

SWAP test circuits were evaluated using two simulation backends: an ideal statevector simulator and a noise-aware device model. The statevector simulator computes circuit outputs under ideal noiseless conditions, whereas the noise-aware backend, FakeWashingtonV2, emulates a superconducting quantum device model incorporating hardware constraints and gate errors.

#### Supplementary Note S3: Worked examples of quantum-state encoding for RNA-seq data

A quantum state corresponding to gene $g$ in the EA model is defined as

$$|\psi_g\rangle = \sum_{t=1}^{T} a_{g,t} e^{i\alpha\phi_{g,t}} |\chi_t\rangle,$$

where $a_{g,t}$, $\phi_{g,t}$, and $|\chi_t\rangle$ denote the amplitude, phase, and the computational basis state representing time point $t$, respectively. The amplitudes satisfy the normalization condition $\sum_{t=1}^{T} |a_{g,t}|^2 = 1$.

For a gene $g$ in the Arabidopsis dataset, which consists of ten time points, the definition can be expanded as

$$|\psi_g\rangle = a_{g,1} e^{i\alpha\phi_{g,1}} |0000\rangle + a_{g,2} e^{i\alpha\phi_{g,2}} |0001\rangle + \cdots + a_{g,10} e^{i\alpha\phi_{g,10}} |1001\rangle$$

using four qubits. Four qubits can represent $2^4 = 16$ basis states. Since the dataset contains only ten time points, the coefficients corresponding to the remaining basis states (e.g., $|1010\rangle, |1011\rangle, \cdots$) are padded to zero.

For a gene $g$ in the wheat dataset, which consists of eight time points, the state can be expanded as

$$|\psi_g\rangle = a_{g,1} e^{i\alpha\phi_{g,1}} |000\rangle + a_{g,2} e^{i\alpha\phi_{g,2}} |001\rangle + \cdots + a_{g,8} e^{i\alpha\phi_{g,8}} |111\rangle$$

using three qubits. In this case, zero-padding was not applied.

#### Supplementary Note S4: Analytical equivalence of fidelity to squared cosine similarity in the expression amplitude model

Consistent with the main text, the quantum state representing gene $g$ is defined as

$$|\psi_g\rangle = \sum_{t=1}^{T} a_{g,t} e^{i\alpha\phi_{g,t}} |\chi_t\rangle.$$

For two genes $g$ and $h$, the fidelity between their corresponding pure states is given by

$$F(g,h) = |\langle\psi_g|\psi_h\rangle|^2 = \left|\sum_{t=1}^{T} a_{g,t} a_{h,t} e^{i\alpha(\phi_{h,t} - \phi_{g,t})}\right|^2.$$

In the EA model, the amplitudes and phases are defined as

$$a_{g,t} = \frac{x_{g,t}}{\sqrt{X_g}}$$

$$\phi_{g,t} = t$$

where $X_g = \sum_{t=1}^{T} |x_{g,t}|^2$.

Since $\phi_{h,t} - \phi_{g,t} = t - t = 0$, the phase factor vanishes, and the fidelity simplifies to

$$F_{EA}(g,h) = \left| \sum_{t=1}^{T} \frac{x_{g,t} x_{h,t}}{\sqrt{X_g X_h}} \right|^2.$$

This expression can be written in vector form as

$$F_{EA}(g,h) = \left( \frac{\boldsymbol{x}_g \cdot \boldsymbol{x}_h}{\|\boldsymbol{x}_g\| \|\boldsymbol{x}_h\|} \right)^2,$$

where $\boldsymbol{x}_g = (x_{g,1}, \ldots, x_{g,T})$.

Because the cosine similarity between two vectors is defined as

$$\cos(\boldsymbol{x}_g, \boldsymbol{x}_h) = \frac{\boldsymbol{x}_g \cdot \boldsymbol{x}_h}{\|\boldsymbol{x}_g\| \|\boldsymbol{x}_h\|},$$

the fidelity in the EA model is exactly the squared cosine similarity between gene expression vectors:

$$F_{\mathrm{EA}}(g,h) = \left(\cos(\boldsymbol{x}_g, \boldsymbol{x}_h)\right)^2.$$

#### Supplementary Note S5: Phase invariance under global sign reversal in the temporal difference phase (TDP) model

Consistent with the main text, in the temporal difference phase (TDP) encoding strategy, the phase of the quantum state at the temporal basis ($|\chi_t\rangle$) is defined as

$$\phi_{g,t} = \Delta z_{g,t} = z_{g,t+1} - z_{g,t},$$

where $z_{g,t}$ denotes the z-score standardized expression magnitude of gene $g$ at time point $t$.

For two genes $g$ and $h$, the phase difference at time index $t$ is given by

$$\phi_{h,t} - \phi_{g,t} = \Delta z_{h,t} - \Delta z_{g,t}.$$

The fidelity between genes $g$ and $h$ is defined as

$$F_{\mathrm{TDP}}(g,h) = \left| \frac{1}{T-1} \sum_{t=1}^{T-1} e^{i\alpha(\phi_{h,t} - \phi_{g,t})} \right|^2 = \left| \frac{1}{T-1} \sum_{t=1}^{T-1} e^{i\alpha(\Delta z_{h,t} - \Delta z_{g,t})} \right|^2.$$

For gene pairs lacking coherent temporal structure, characterized by stochastic fluctuations around a mean, the phase differences $\Delta z_{h,t} - \Delta z_{g,t}$ vary irregularly across time. As a result, the complex exponential terms $e^{i\alpha(\Delta z_{h,t} - \Delta z_{g,t})}$ become effectively uncorrelated, leading to destructive interference in the summation. Consequently, the resulting fidelity tends to take low values.

In comparison, consider gene pairs exhibiting opposing monotonic trends, such as one increasing and the other decreasing over time. In such cases, the temporal differences approximately satisfy

$$\Delta z_{g,t} \approx -\Delta z_{h,t} \quad \text{for all } t.$$

The phase difference then becomes

$$\phi_{h,t} - \phi_{g,t} \approx \Delta z_{h,t} - (-\Delta z_{h,t}) = 2\Delta z_{h,t},$$

and the corresponding phase factors evolve smoothly over time.

When the temporal changes are gradual, the phase differences $\phi_{h,t} - \phi_{g,t}$ also vary smoothly rather than randomly. This structured evolution prevents complete cancellation in the complex summation and induces partial constructive interference. As a result, even though the expression trajectories exhibit globally opposite trends, the fidelity can attain relatively high values.

#### Supplementary Note S6: Insensitivity to initial temporal discrepancies in the integrated difference phase (IDP) model

Consistent with the main text, in the integrated difference phase (IDP) encoding strategy, the phase of the quantum state at temporal basis ($|\chi_t\rangle$) is defined as

$$\phi_{g,t} = \sum_{k=1}^{t} \omega_{g,k} = \sum_{k=1}^{t} \frac{\Delta z_{g,k}}{\sigma_\Delta},$$

where $\Delta z_{g,t} = z_{g,t+1} - z_{g,t}$ and $z_{g,k}$ denotes the z-score standardized expression magnitude of gene $g$ at time point $t$.

Because the cumulative sum of successive differences forms a telescoping sum,

$$\sum_{k=1}^{t} \Delta z_{g,k} = z_{g,t+1} - z_{g,1},$$

the phase can be rewritten as

$$\phi_{g,t} = \frac{1}{\sigma_\Delta}(z_{g,t+1} - z_{g,1}).$$

Thus, the IDP encoding represents the cumulative deviation of expression from the initial time point.

For two genes $g$ and $h$, the phase difference at time index $t$ is given by

$$\phi_{h,t} - \phi_{g,t} = \frac{1}{\sigma_\Delta}\left((z_{h,t+1} - z_{h,1}) - (z_{g,t+1} - z_{g,1})\right).$$

The fidelity between genes $g$ and $h$ is defined as

$$F_{\text{IDP}} = \left| \frac{1}{T-1} \sum_{t=1}^{T-1} e^{i\alpha(\phi_{h,t} - \phi_{g,t})} \right|^2.$$

From this formulation, it follows that the magnitude of fidelity is governed by the temporal behavior of the phase differences. When the phase differences $\phi_{h,t} - \phi_{g,t}$ vary irregularly across time, the complex exponential terms become effectively uncorrelated, resulting in destructive interference and low fidelity values.

In contrast, high fidelity arises when the phase differences remain approximately constant over time. From the expression above, this condition is satisfied when the difference in expression between the two genes is temporally stable, i.e.,

$$z_{h,t} - z_{g,t} \approx \delta$$

for all $t$, where $\delta$ is a constant offset. In this case,

$$\phi_{h,t} - \phi_{g,t} = \frac{1}{\sigma_\Delta}\Big(\big(z_{g,t} - z_{h,1}\big) - \delta\Big),$$

which is independent of $t$. Under this condition, the phase factors $e^{i\alpha(\phi_{h,t}-\phi_{g,t})}$ become nearly identical across time points, leading to constructive interference in the summation. Consequently, the fidelity attains relatively high values.

Notably, this condition can still be satisfied even when the initial temporal changes differ substantially between genes. For example, opposing transitions at early time points may introduce a constant offset in the cumulative phase. However, if subsequent expression trajectories evolve in a parallel manner such that the difference $z_{h,t} - z_{g,t}$ remains approximately constant for $t \geq 2$, the phase difference remains approximately constant over time. In this case, the magnitude of the phase difference is not critical; rather, its temporal stability ensures that the corresponding phase factors remain coherent across time, leading to relatively high-fidelity values.

### Supplementary Tables

#### Supplementary Table S1

**Distribution statistics for fidelity values and Pearson correlation coefficients.** For each dataset and model or reference measure, the table reports the mean, standard deviation, minimum, first quartile (Q1), median (Q2), third quartile (Q3), maximum, and interquartile range (IQR). $PCC_e$ and $PCC_d$ denote Pearson correlation coefficients computed from expression profiles and temporal difference profiles, respectively.

#### Supplementary Table S2

**Mean absolute errors between encoding models and reference measures.** Values represent mean absolute errors between the pairwise similarity values obtained from the models or reference measures shown in the corresponding row and column. $PCC_e$ and $PCC_d$ denote Pearson correlation coefficients computed from expression profiles and temporal difference profiles, respectively.

#### Supplementary Table S3

**Discordant gene pairs identified using threshold criteria. Pair counts were computed from the full pairwise similarity matrices for each dataset.** Discordant pairs were defined as gene pairs showing high fidelity despite strong negative Pearson correlation based on expression profiles. $PCC_e$ denotes the Pearson correlation coefficient computed from expression profiles.

#### Supplementary Table S4

**Agreement between communities detected from averaged expression and replicate-resampled fidelity matrices.** Networks were reconstructed from replicate-resampled mean fidelity matrices computed across 1,000 sample-level resampling iterations using the same parameters selected in the main averaged-expression analysis. The table reports the number of input genes, the number of genes assigned to retained communities in each analysis, the Jaccard index of assigned gene sets, adjusted Rand index (ARI), adjusted mutual information (AMI), normalized mutual information (NMI), and variation of information (VI). Metrics are reported both for all input genes, with unassigned genes treated as a separate category, and for genes assigned to communities in both analyses.

### Supplementary Figures

#### Supplementary Figure S1

**Pairwise relationships of similarity values across encoding strategies.** Pairwise comparisons of fidelity values across encoding strategies using 100,000 randomly sampled gene pairs for (A) Arabidopsis, (B) wheat, (C), soybean, (D) human, and (E) mouse datasets. Each panel shows the relationship between the pairwise similarity values obtained from the corresponding models or reference measures.

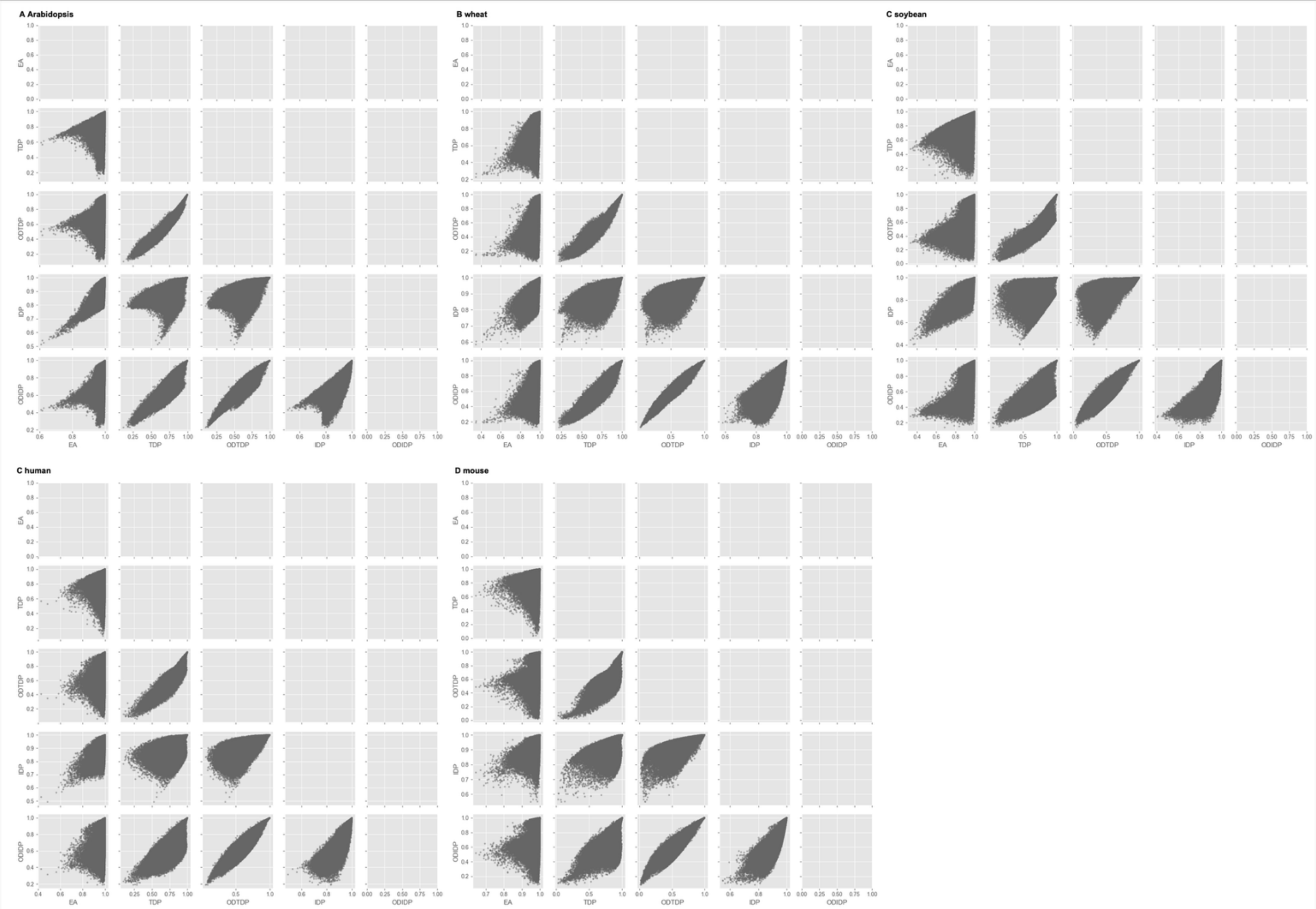

#### Supplementary Figure S2

**Examples of high fidelity with low Pearson correlation under phase-based encodings.** Representative gene pairs from the wheat dataset are shown for cases in which fidelity was high despite low Pearson correlation. In each panel, the upper plots show variance-stabilized expression values across time points for the two genes. The Pearson correlation coefficient and fidelity value for each gene pair are shown below the corresponding expression plots. (A) Two representative gene pairs from the TDP model. (B) Two representative gene pairs from the IDP model.

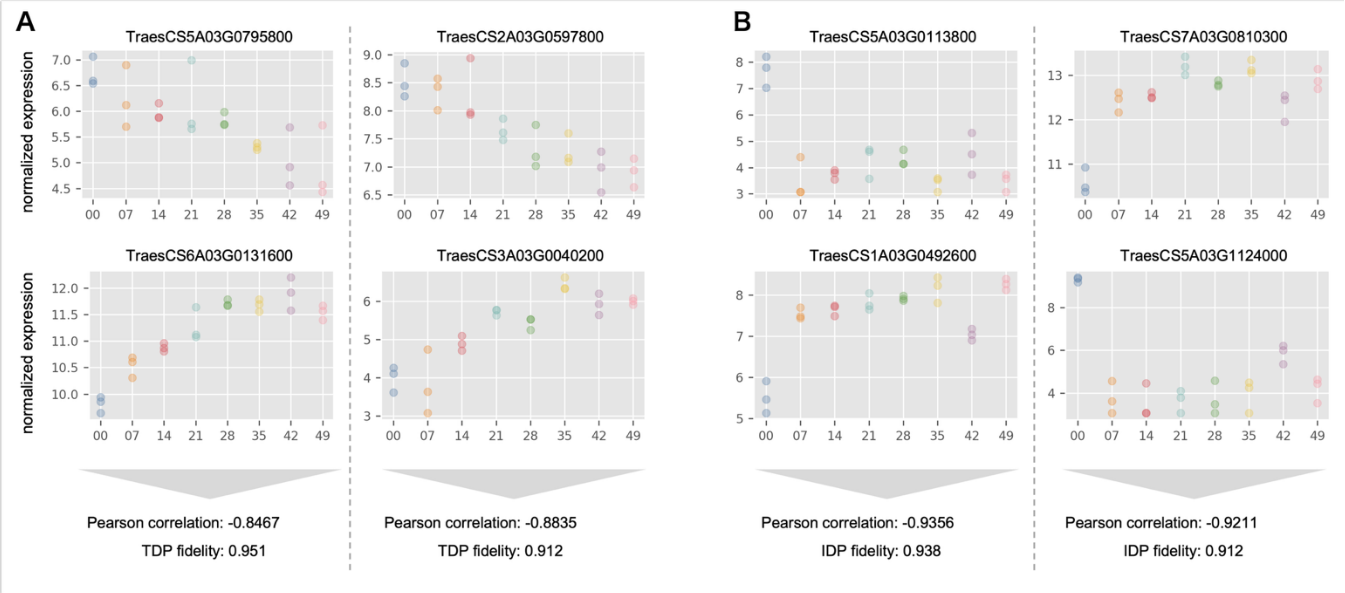

#### Supplementary Figure S3

**Temporal expression patterns within detected communities in the Arabidopsis dataset.** Temporal expression profiles of genes in communities detected by the fidelity-based models, WGCNA, and Mfuzz. Fidelity based communities were detected using the default phase scaling parameter. Each line represents one gene, and the number of genes in each community is shown in the corresponding panel title.

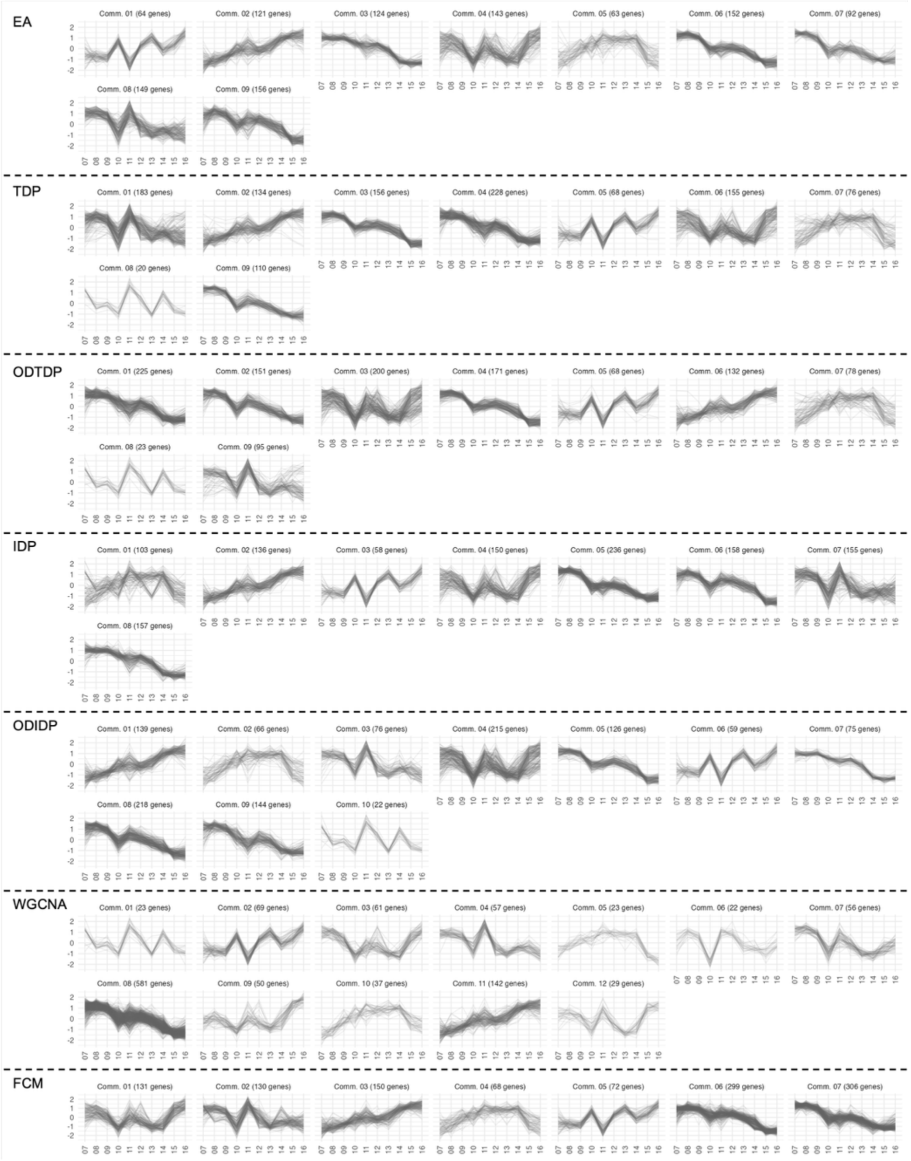

#### Supplementary Figure S4

**Temporal expression patterns within detected communities in the wheat dataset.** Temporal expression profiles of genes in communities detected by the fidelity-based models, WGCNA, and Mfuzz. Fidelity based communities were detected using the default phase scaling parameter. Each line represents one gene, and the number of genes in each community is shown in the corresponding panel title.

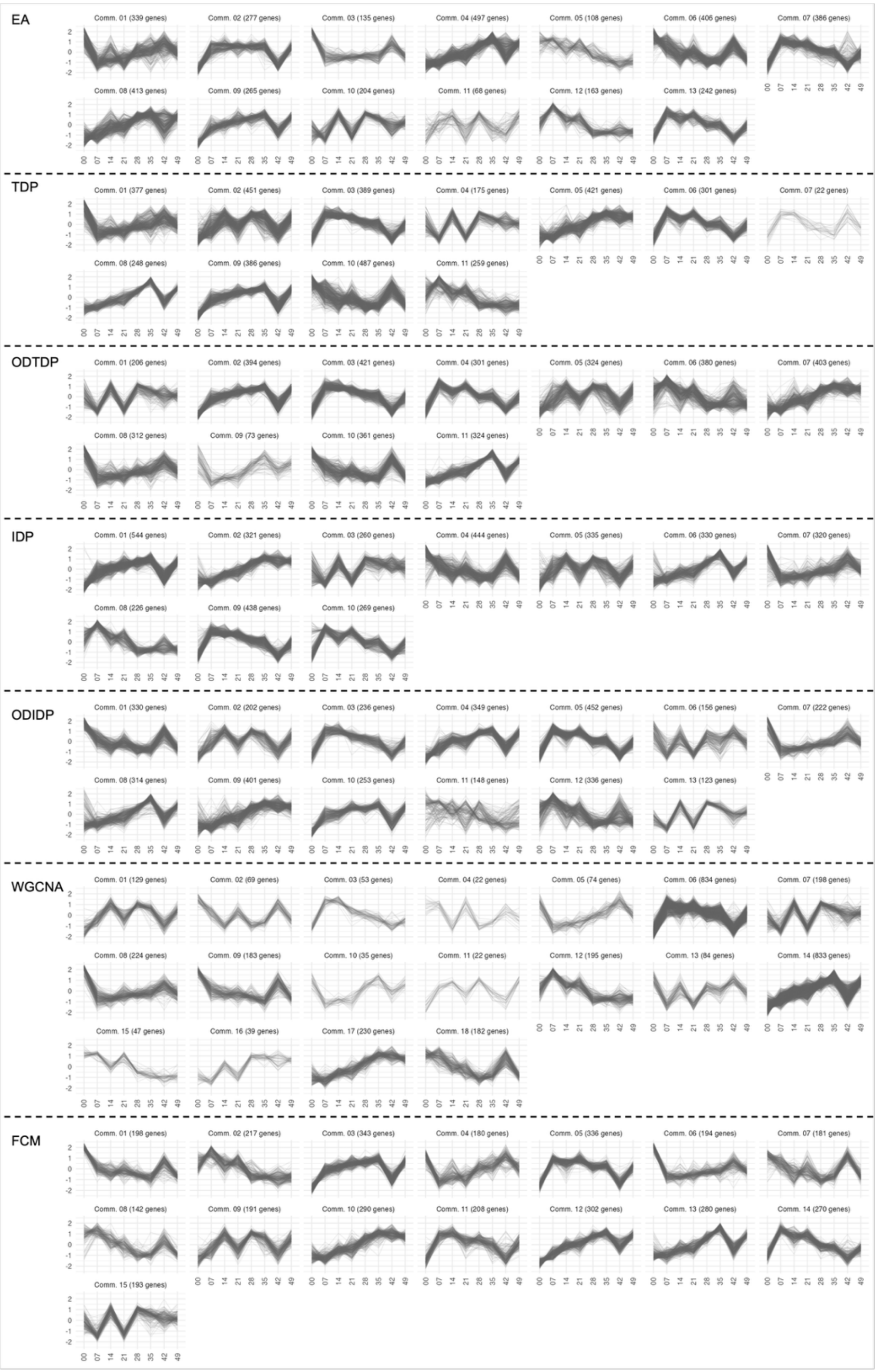

#### Supplementary Figure S5

**Temporal expression patterns within detected communities in the soybean dataset.** Temporal expression profiles of genes in communities detected by the fidelity-based models, WGCNA, and Mfuzz. Fidelity based communities were detected using the default phase scaling parameter. Each line represents one gene, and the number of genes in each community is shown in the corresponding panel title.

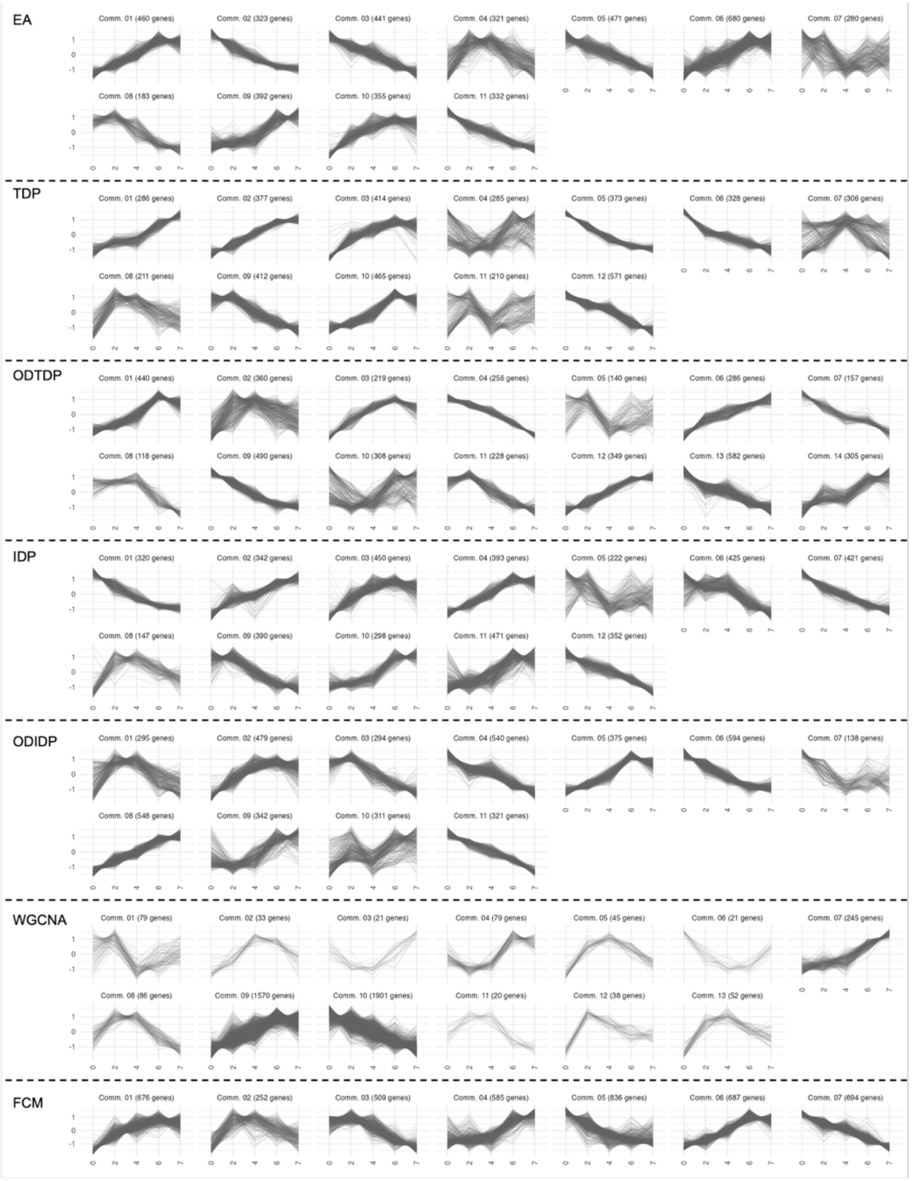

#### Supplementary Figure S6

**Temporal expression patterns within detected communities in the human dataset.** Temporal expression profiles of genes in communities detected by the fidelity-based models, WGCNA, and Mfuzz. Fidelity based communities were detected using the default phase scaling parameter. Each line represents one gene, and the number of genes in each community is shown in the corresponding panel title.

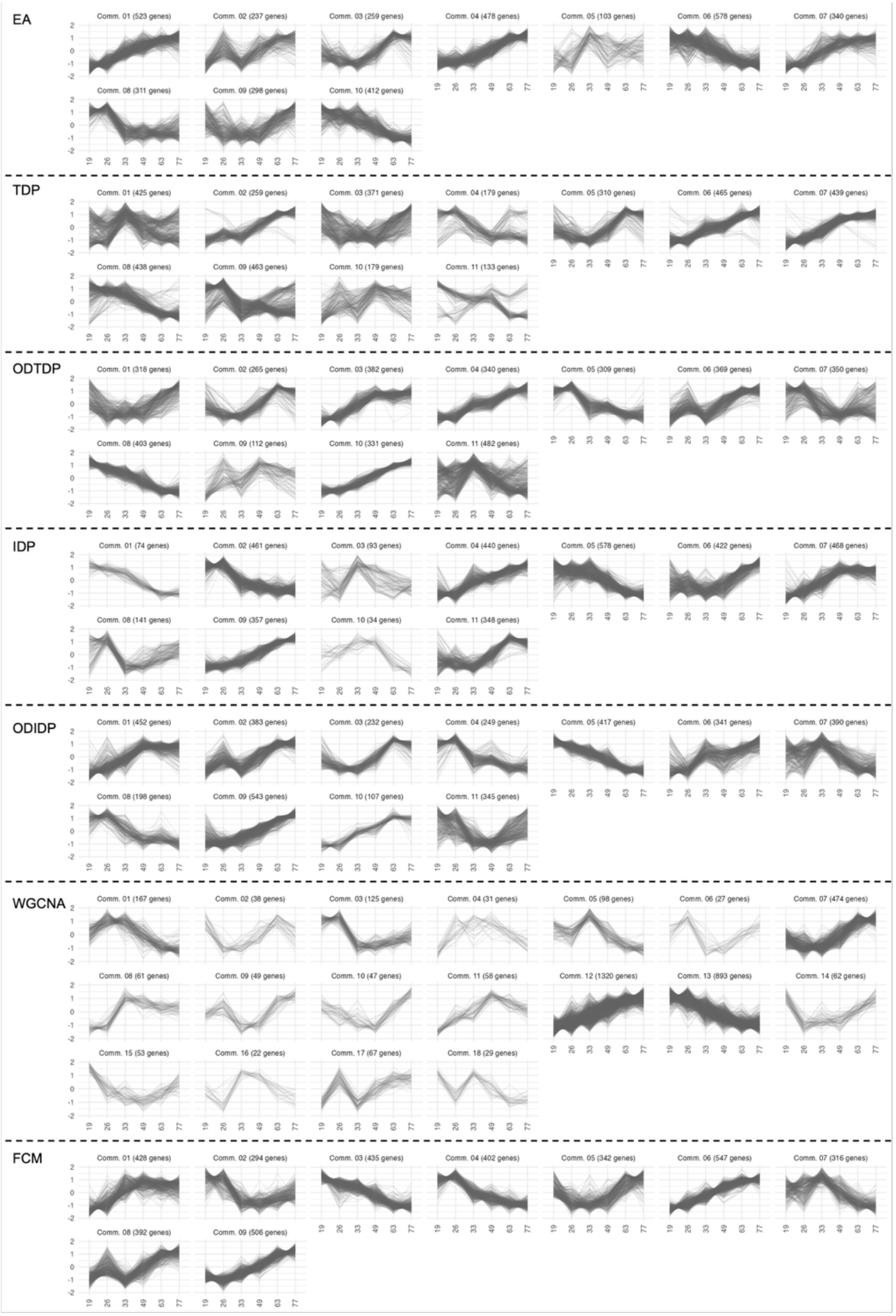

#### Supplementary Figure S7

**Temporal expression patterns within detected communities in the mouse dataset.** Temporal expression profiles of genes in communities detected by the fidelity-based models, WGCNA, and Mfuzz. Fidelity based communities were detected using the default phase scaling parameter. Each line represents one gene, and the number of genes in each community is shown in the corresponding panel title.

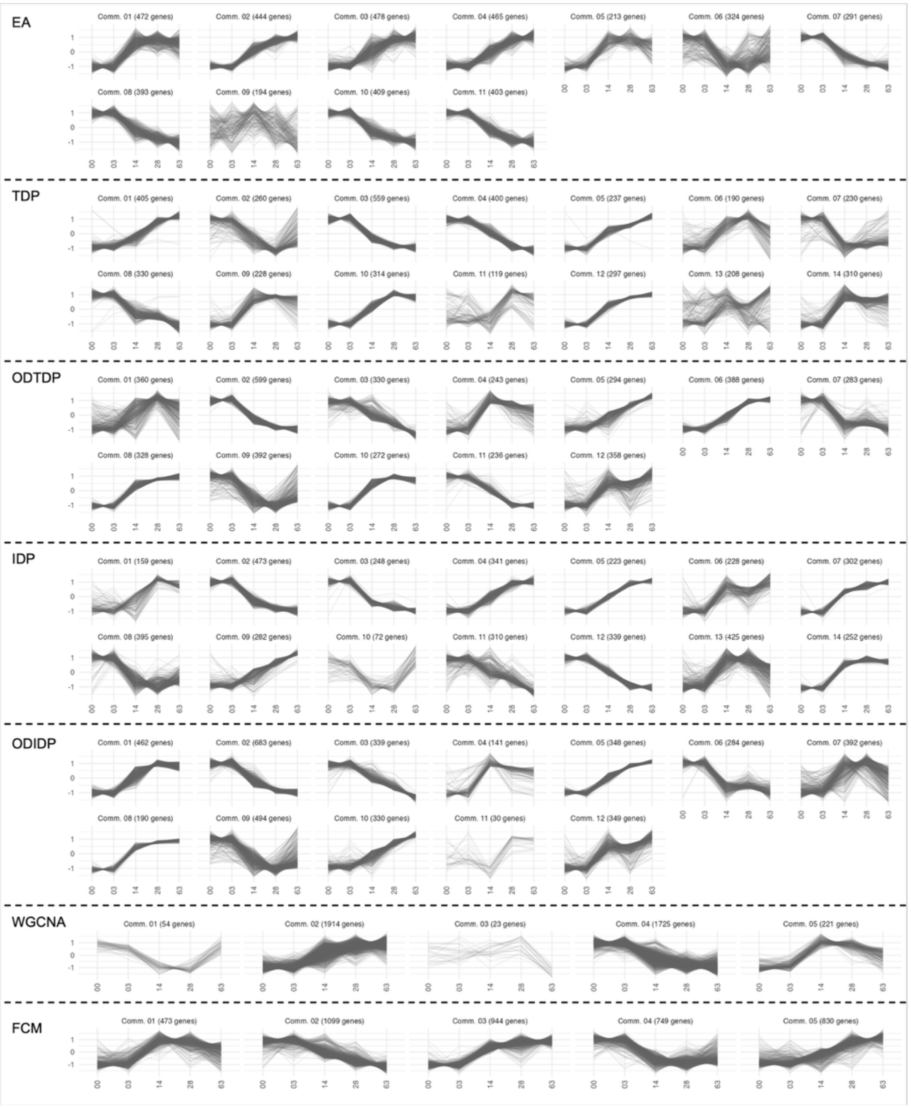

#### Supplementary Figure S8

**Relationship between fidelity values computed from expression values averaged across replicates and from resampled expression matrices.** Scatter plots compare pairwise fidelity values computed from expression values averaged across biological replicates with resampled mean fidelity values computed from 1,000 resampling iterations. Each point represents a gene pair included in the comparison. The dashed line indicates equality between the two fidelity estimates. Values in each panel indicate the Pearson correlation coefficient (PCC) and mean absolute error (MAE) between the two sets of fidelity values. Each point represents one of 100,000 randomly sampled gene pairs.

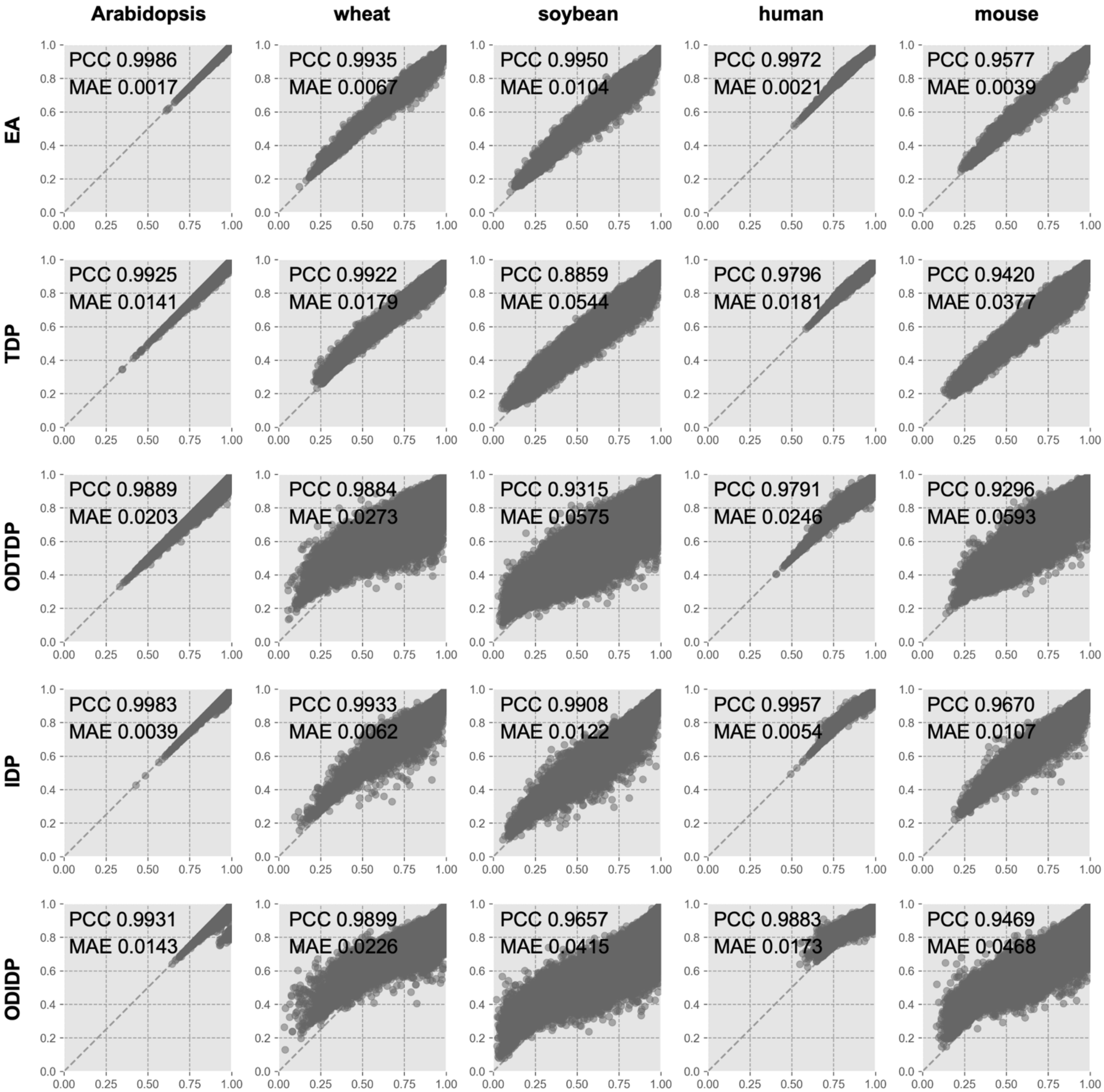

#### Supplementary Figure S9

**Temporal expression profiles of communities detected from replicate-resampled mean fidelity matrices.** Heatmaps show the mean standardized expression profiles of communities reconstructed from replicate-resampled mean fidelity matrices. The layout follows that of Fig. 2 to facilitate comparison with communities detected from averaged expression profiles across replicates, where (A) Arabidopsis, (B) wheat, (C) soybean, (D) human, and (E) mouse datasets. Replicate-resampled mean fidelity matrices were computed from 1,000 sample-level resampling iterations, and networks were reconstructed using the same parameters selected in the main averaged-expression analysis without re-optimization.

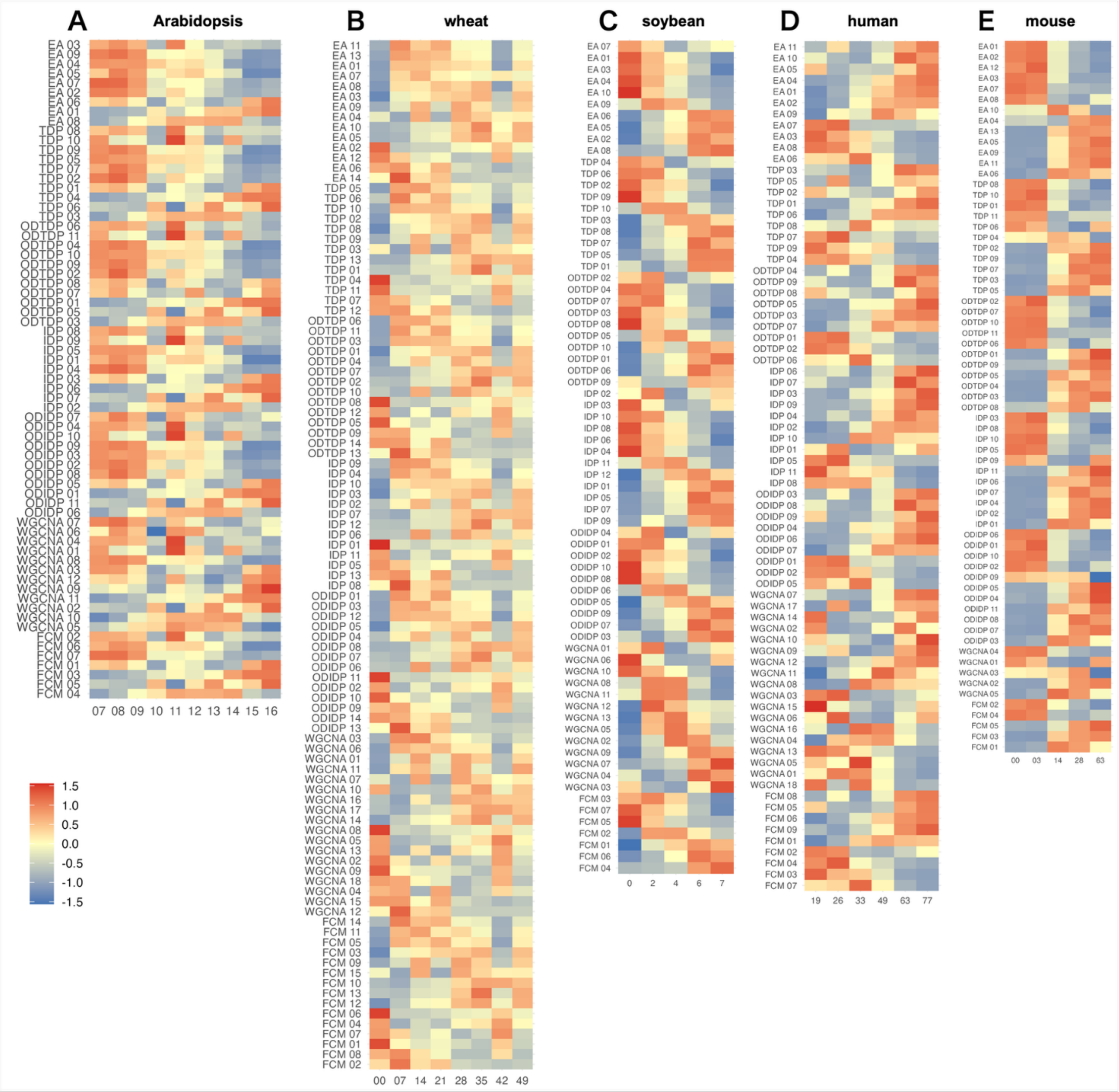

#### Supplementary Figure S10

**Fidelity distributions across encoding strategies and phase scaling values.** Histograms of pairwise fidelity values computed from 100,000 randomly sampled gene pairs across encoding strategies and phase scaling values for (A) Arabidopsis, (B) wheat, (C), soybean, (D) human, and (E) mouse datasets.

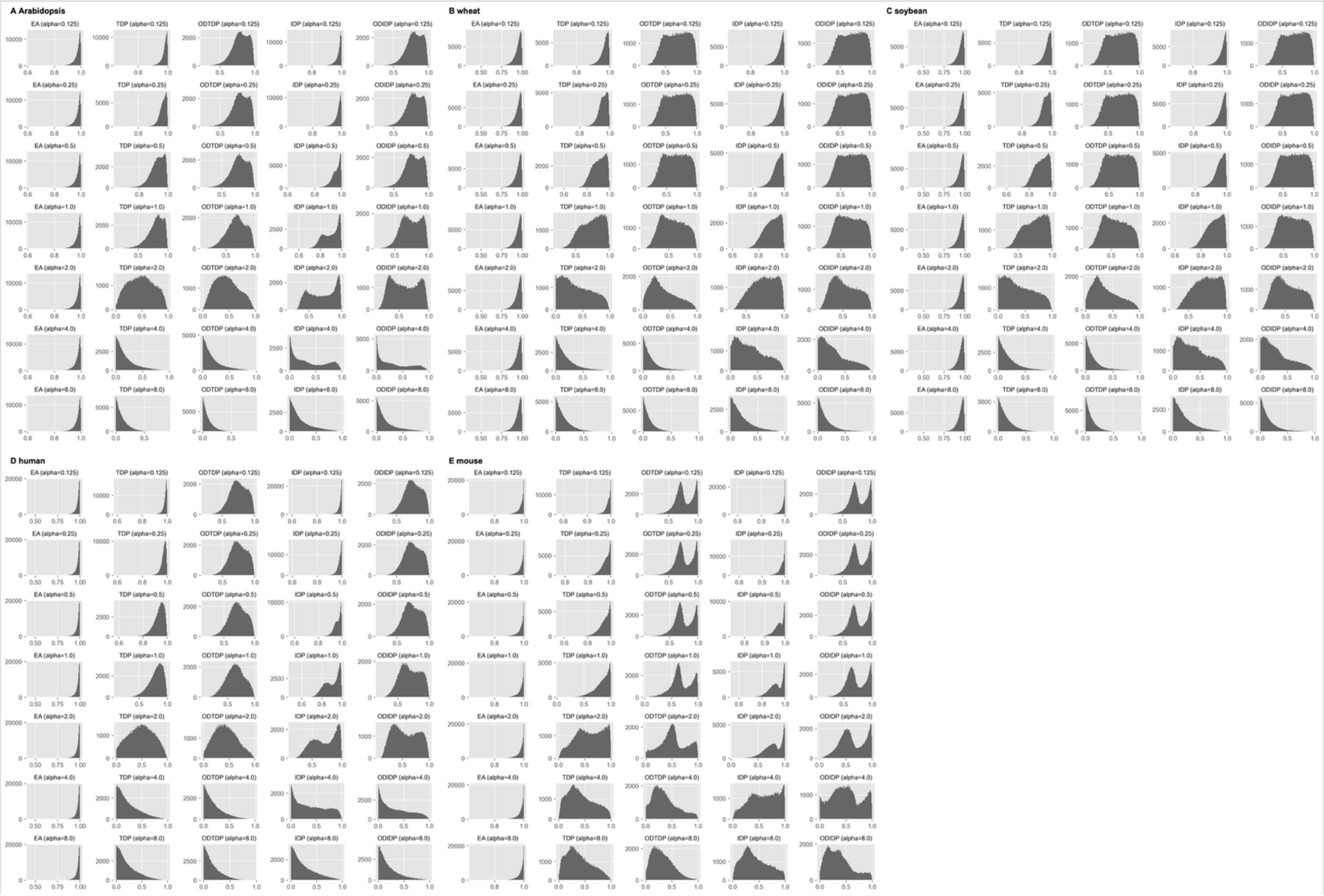

#### Supplementary Figure S11

**Temporal expression patterns within detected communities in the human dataset under larger phase scales.** Temporal expression profiles of genes in communities detected by the fidelity-based models under the selected phase scaling condition. Each line represents one gene, and the number of genes in each community is shown in the corresponding panel.

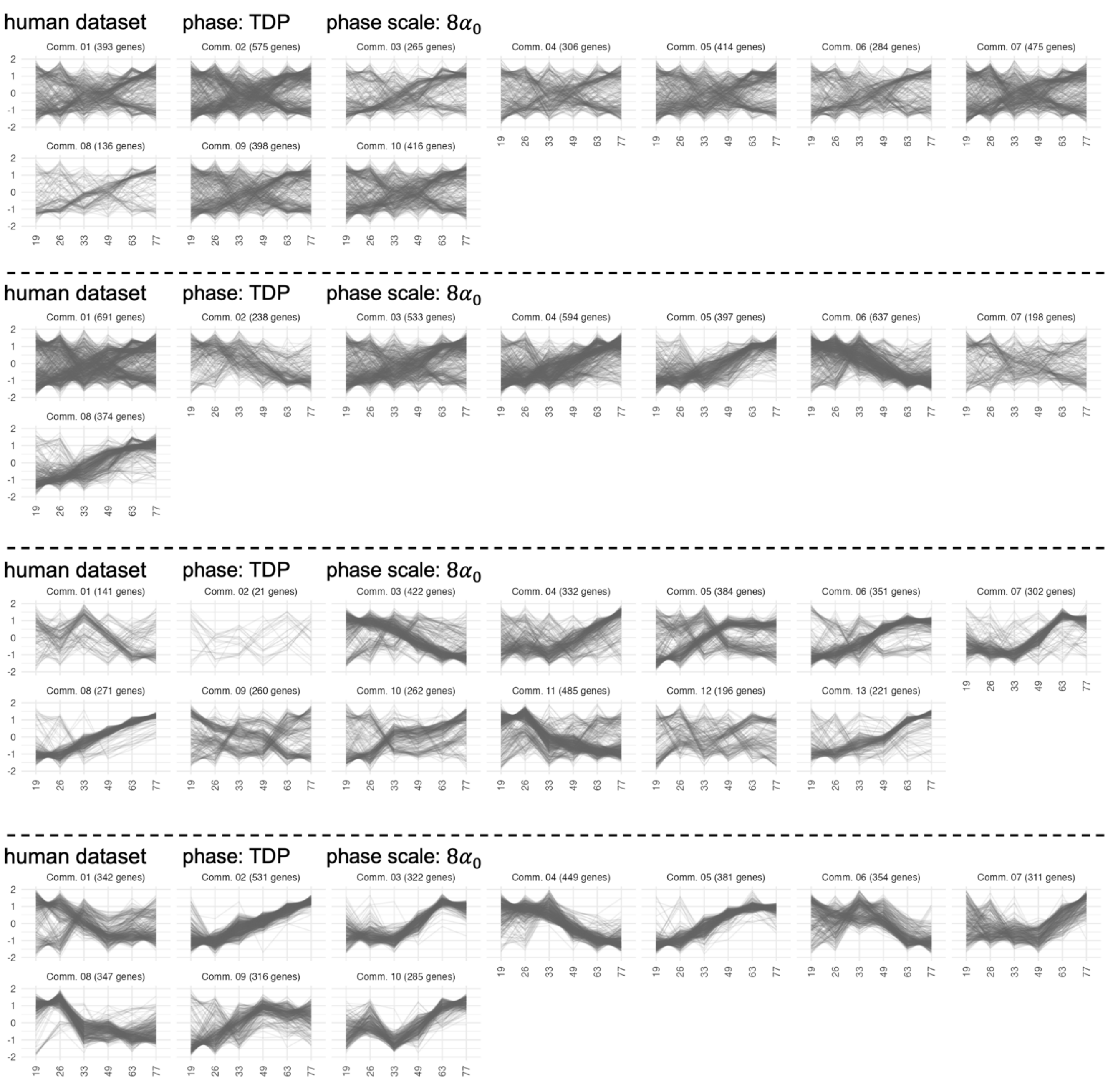
